# Tmbim5 and Slc8b1 cooperate in tissue-specific mitochondrial calcium regulation in zebrafish

**DOI:** 10.1101/2024.12.03.626543

**Authors:** Iga Wasilewska, Łukasz Majewski, Dobrochna Adamek-Urbańska, Sofiia Baranykova, Matylda Macias, Aleksandra Szybińska, Axel Methner

## Abstract

Mitochondrial calcium homeostasis involves coordinated uptake via the mitochondrial calcium uniporter (MCU) and efflux through sodium-dependent NCLX (encoded by *SLC8B1*). Mild *Mcu* knockout phenotypes suggest additional transport mechanisms. We investigated TMBIM5, a proposed bidirectional mitochondrial calcium/proton transporter, by generating zebrafish lacking *tmbim5*, *slc8b1*, plus *tmbim5/mcu* and *tmbim5/slc8b1* double knockouts. Tmbim5-deficient fish exhibited growth impairment, muscle atrophy, and increased brain cell death. *tmbim5/mcu* double knockouts showed no additive effects, arguing against Tmbim5 functioning as an independent calcium uptake pathway. However, *tmbim5/slc8b1* double knockouts showed disrupted mitochondrial calcium handling with reduced uptake and efflux. Remarkably, brain phenotypes were rescued while muscle dysfunction was exacerbated in double mutants, corresponding to restored mitochondrial membrane potential in Tmbim5-deficient brain tissue and exacerbated decreased calcium levels in double knockout muscle. We also found broad downregulation of mitochondrial calcium transport proteins and decreased mitochondrial DNA content in double knockout brain tissue, indicating reduced mitochondrial mass and a potential beneficial glycolytic shift in energy metabolism. These findings demonstrate that TMBIM5 functions as a calcium efflux pathway cooperating with NCLX in a tissue-specific manner.

## Introduction

Mitochondria generate ATP by harnessing proton and electron gradients established by the electron transport system. The negative mitochondrial membrane potential across the inner mitochondrial membrane also serves as the driving force for uptake of the universal cellular messenger calcium (Ca^2+^). Increased mitochondrial Ca^2+^ levels enhance oxidative phosphorylation, leading to increased ATP production. Additionally, mitochondria function as cellular Ca^2+^ sinks, making mitochondrial Ca^2+^ uptake crucial for cellular Ca^2+^ signaling. Therefore, mitochondrial Ca^2+^ homeostasis represents a finely tuned process involving the coordinated action of Ca^2+^ transporters, regulatory proteins, and interactions with the endoplasmic reticulum (ER).

The mechanisms governing mitochondrial Ca^2+^ transport remain incompletely understood. The mitochondrial Ca^2+^ channel mediating inward Ca^2+^ flux is the mitochondrial Ca^2+^ uniporter (MCU) ^1,2^, which forms part of a large protein complex containing several MCU-interacting and regulatory proteins including MICU1 (mitochondrial uptake 1) ^3^, MICU2 ^4^, EMRE (essential MCU regulator) ^5^, and MCUR1 (mitochondrial Ca^2+^ uniporter regulator 1) ^6,7^. Despite its central role, *Mcu* knockout (KO) produces remarkably mild phenotypes: in mice, *Mcu* deletion causes mild phenotypes in a CD1 background ^8^ but embryonic lethality in C57Bl/6 ^9^. In zebrafish, *mcu* knockdown or knockout had minimal effects on survival, activity, viability, fertility, morphology, or swimming behavior, although it abolished mitochondrial Ca^2+^ uptake similar to the well-characterized MCU inhibitor ruthenium red (RuR) ^10–13^. *In vivo* studies in zebrafish photoreceptors suggest alternative Ca^2+^ entry pathways can compensate for Mcu loss ^14^.

One disputed candidate for alternative Ca^2+^ transport is Leucine Zipper And EF-Hand Containing Transmembrane Protein 1 (LETM1), identified by genome-wide screening as a RuR-sensitive mitochondrial Ca^2+^/H^+^-antiporter ^15^. Originally, LETM1 was described as part of the mitochondrial K^+^/H^+^ exchanger ^16^. For Ca^2+^ efflux, the primary mediator is the Na^+^/Ca^2+^ exchanger NCLX ^17^, whose knockout causes embryonic lethality in mice ^18^. Recent evidence suggests that Transmembrane Protein 65 (TMEM65) is also involved in Na^+^-dependent Ca^2+^ efflux from mitochondria ^19,20^ and may act as a positive regulator of NCLX ^21^. Another mitochondrial Ca^2+^ extrusion system is the mitochondrial permeability transition pore (mPTP), a non-selective high-conductance pore of debated identity ^22,23^. High mitochondrial Ca^2+^ concentrations trigger mPTP opening, which when sustained causes mitochondrial swelling, breakdown of mitochondrial membrane potential, and eventually cell death (reviewed in ^24^).

Recently, transmembrane BAX inhibitor motif containing protein 5 (TMBIM5) has emerged as a novel candidate for mitochondrial Ca^2+^ transport. TMBIM5 was initially linked to regulation of mitochondrial morphology, as its loss resulted in mitochondrial fragmentation and disrupted cristae structure ^25–28^. Loss of TMBIM5 also increased cellular sensitivity to apoptosis ^25,26,29^. *In vivo*, mice with a specific alteration (D326R) that presumably disrupts the channel pore of TMBIM5 ^30^ and leads to protein degradation ^27^ exhibit higher embryonic mortality and develop skeletal myopathy characterized by reduced muscle strength. These observations correlated with several mitochondrial abnormalities, most dramatically in skeletal muscle mitochondria, including disrupted cristae structure, premature mPTP opening, decreased mitochondrial Ca^2+^ uptake capacity, and mitochondrial swelling ^27^.

Two independent studies demonstrated that TMBIM5, when reconstituted in liposomes, can function as a Ca^2+^ transporter in the presence of a pH gradient ^28,29^. Inhibiting NCLX-mediated mitochondrial Ca^2+^ efflux with CGP-37157 ^28^ or *NCLX* knockdown ^29^ resulted in diminished mitochondrial Ca^2+^ retention capacity in *TMBIM5* KO human embryonic kidney (HEK) cells and reduced RuR-induced mitochondrial Ca^2+^ efflux ^28^. *TMBIM5* KO cells also accumulated more matrix Ca^2+^ upon ER Ca^2+^ release mediated by inhibition of the sarco/endoplasmic reticulum ATPase (SERCA) with thapsigargin ^29^ or stimulation of inositol trisphosphate receptor-mediated Ca^2+^ release with ATP or carbachol ^28^. Restricting mitochondrial Ca^2+^ uptake by concurrent *MCU* knockout rescued elevated apoptosis induced by staurosporine in *TMBIM5*^-/-^ cells ^29^. Collectively, these data suggest that TMBIM5 can mediate Ca^2+^ extrusion from mitochondria and that its function at least partially overlaps with NCLX.

In contrast, own previous work found that TMBIM5 overexpression in HEK cells enhanced mitochondrial Ca^2+^ uptake following ER Ca^2+^ release ^27^, suggesting that TMBIM5 may operate bidirectionally under certain conditions. This study also demonstrated that under steady-state conditions, loss of TMBIM5 increases potassium and reduces proton levels in the mitochondrial matrix, suggesting a broader role in mitochondrial ion exchange ^27^, consistent with a potential interaction between TMBIM5 and LETM1 ^28^. However, the physiological significance of these mechanisms *in vivo*, particularly the crosstalk between TMBIM5, MCU, and NCLX, remains unclear.

Due to their translucency, zebrafish represent an ideal model to study mitochondrial Ca^2+^ dynamics *in vivo* ^31,32^. In the present study, we generated *tmbim5* and *slc8b1* KO fish using CRISPR/Cas9-mediated gene editing ^33^ and observed that Tmbim5 loss results in a muscle phenotype associated with delayed hatching, reduced size, and increased brain cell death. To investigate potential interactions with known Ca^2+^ transport systems, we generated *tmbim5*/*mcu* and *tmbim5*/*slc8b1* double knockouts. Both lines remained viable without major phenotypes; however, *tmbim5*/*slc8b1* double knockouts showed tissue-specific rescue or aggravation of phenotypes associated with Tmbim5 loss of function. These findings suggest that mitochondrial Ca^2+^ transport systems operate in a highly tissue-specific manner in fish.

## Results

### Loss of Tmbim5 impairs zebrafish development and muscle function

The zebrafish genome contains only one *TMBIM5* orthologue which is expressed as early as six hours post fertilization (hpf) and plateaus after 24 hours (Fig. 1A). To study the effect of Tmbim5 deficiency, we generated mutant fish by introducing a deletion in exon 4 using CRISPR/Cas9 genome editing ^33^. This resulted in a profound decrease in *tmbim5* mRNA (Fig. 1B). Mutants (*tmbim5*^−/−^) reached adulthood and the Mendelian distribution was unchanged (Fig. 1C) indicating no increased embryonic lethality (Fig. 1D).

**Figure 1.**
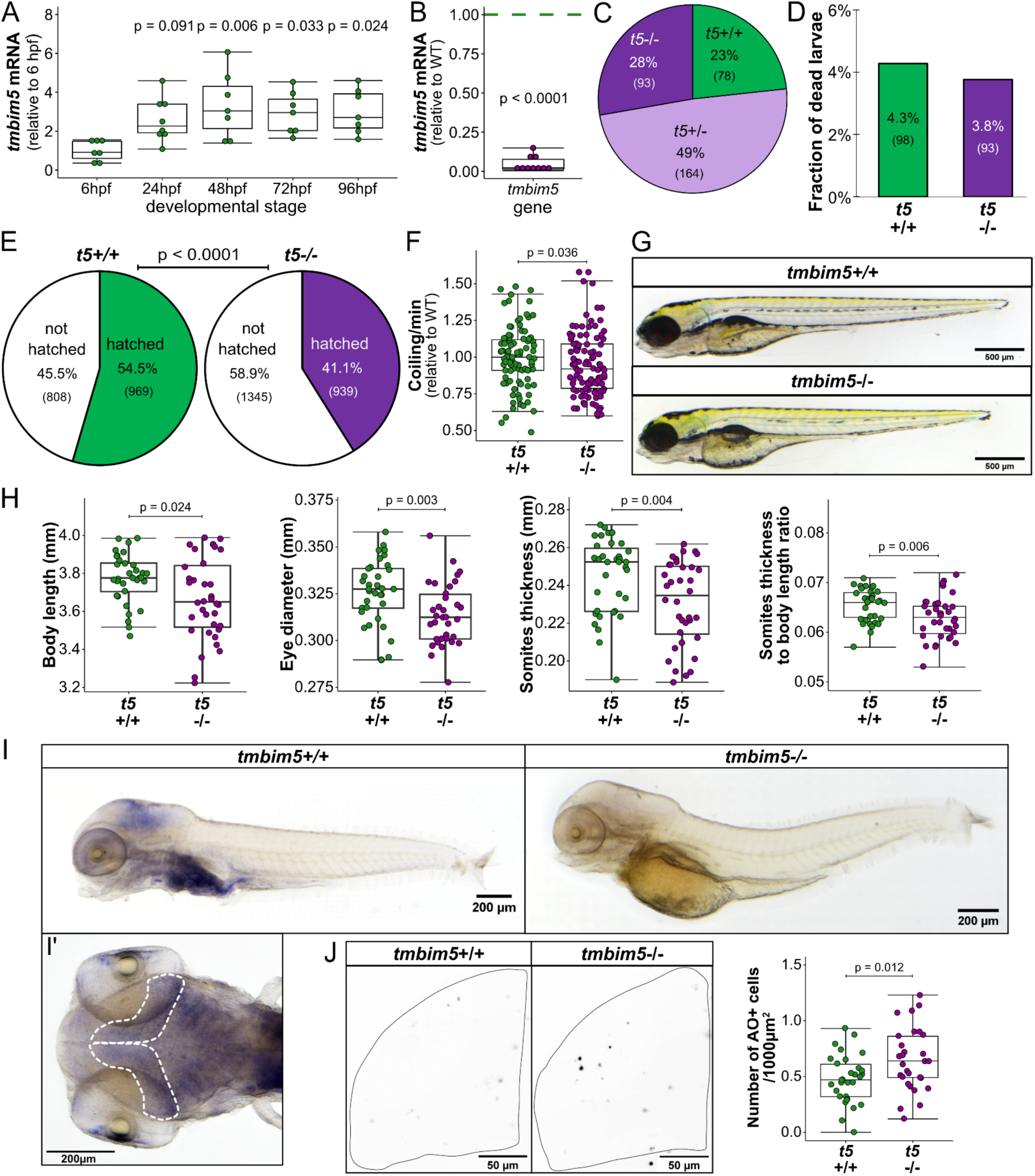
Loss of Tmbim5 impairs zebrafish development and muscle function. A) *tmbim5* mRNA levels at different developmental stages, measured using quantitative PCR (qPCR). Expression was normalized to 6 hours post-fertilization (hpf), with *18S* used as a reference gene. Data are presented as box-and-whisker plots, where the box represents the 25th–75th percentile, and the whiskers indicate the minimum and maximum values. Each dot represents an independent biological replicate (*n* = 7–8; each RNA sample was isolated from 30 embryos/larvae). Number of experiments = 4. Statistical analysis: one-way ANOVA followed by Tukey’s HSD test; *p*-values indicate comparisons with 6 hpf. B) Decreased *tmbim5* mRNA levels in 5 days post-fertilization (dpf) *tmbim5*^-/-^ larvae, quantified using qPCR and normalized to WT levels. *rpl13a* and *ef1a* were used as reference genes. Data are shown as box-and-whisker plots. Each dot represents an independent biological replicate (*n* = 10; each RNA sample was isolated from 30 larvae). Number of experiments = 5. Statistical analysis: one-sample *t*-test. C) Genotype distribution of offspring from *tmbim5* (*t5*)^+/-^ zebrafish breeding does not deviate from expected Mendelian ratios (*n* indicated in brackets). Number of experiments = 4. Statistical analysis: Chi-square test. D) Survival of *tmbim5*^-/-^ larvae up to 5 dpf is not significantly affected. *n* is indicated in brackets. Number of experiments = 7. Statistical analysis: Chi-square test. E) Reduced hatching efficiency of *tmbim5*^-/-^ larvae. *n* is indicated in brackets. Number of experiments = 6. Statistical analysis: Chi-square test. F) Decreased coiling activity of *tmbim5*^-/-^ embryos at 1 dpf. The mean number of coiling movements per minute, normalized to WT from the same experiment, is shown as box-and-whisker plots. Each dot represents an individual embryo (*n* = 94–104). Number of experiments = 6. Statistical analysis: one-sample Wilcoxon test. G) Normal morphology of *tmbim5*^-/-^ larvae at 5 dpf. Representative images of WT (*tmbim5*^+/+^) and *tmbim5*^-/-^ larvae are shown. H) Reduced size of *tmbim5*^-/-^ larvae. Morphometric analysis results are presented as box-and-whisker plots, with each dot representing an individual larva (*n* = 34–36). Number of experiments = 3. Statistical analysis: Mann–Whitney test (for body length and somite thickness) or *t*-test. I) *tmbim5* transcript detected using whole-mount in situ hybridization in 4 dpf wild-type (WT) larvae, with a complete loss of expression in *tmbim5*^-/-^ larvae. I’ Dorsal view of the optic tectum region outlined with a dashed line. Number of experiments = 4. J) Increased cell death in the optic tectum of *tmbim5^-/-^* larvae detected using Acridine Orange (AO) staining. Representative images of WT and *tmbim5*^-/-^ larvae (black dots indicate dying cells with high AO signals). Quantification results are presented as box-and-whisker plots, with each dot representing an individual larva (*n* = 29–30, number of experiments = 3). Statistical analysis: *t*-test.

Mutant larvae took longer to hatch than wild-type (WT, *tmbim5*^+/+^) controls (Fig. 1E) and this correlated with reduced tail coiling (Fig. 1F), spontaneous movements of the tail that enable the embryo to escape from its chorion ^34^. This suggested a possible reduction in muscle strength as also observed in mice with a channel pore mutation in TMBIM5 ^27^. Despite that, mutant larvae displayed unchanged spontaneous locomotor activity in an open field test (Fig. S1A) as well as after stimulating mobility in a visual-motor response test (Fig. S1B).

Tmbim5-deficient larvae were morphologically normal (Fig. 1G) but shorter, had a smaller eye diameter and thinner somites – segments of trunk and tail of larvae (Fig. 1H). This reduction in somite thickness was not solely attributable to the overall reduction in whole-body size, as the ratio of somite thickness to whole-body length was also decreased in *tmbim5*^−/−^ larvae (Fig. 1H). Using whole-mount *in situ* hybridization we found that *tmbim5* is strongly expressed in the digestive system, gills, eyes and head with an especially high signal in the area of the midbrain of zebrafish larvae (Fig. 1I). Importantly no staining was observed in *tmbim5*^−/−^ larvae proving the specificity of the riboprobe. Based on the strong expression of *tmbim5* in the brain, we quantified cell death in this tissue by staining the brain of *tmbim5*^−/−^ larvae with Acridine Orange. This demonstrated increased cell death (Fig. 1J) caused by Tmbim5 deficiency. Together these results suggest a role for Tmbim5 in body size, muscle strength and cell viability in the nervous system.

### Tmbim5 deficiency causes muscle atrophy with preferential effects on slow-twitch fibers

We next studied the effects of Tmbim5 deficiency in adult zebrafish to determine whether the developmental phenotypes persist into adulthood. Similar to larvae, *tmbim5*^−/−^ adult fish had a normal morphology, a tendency to be smaller and a significant reduction in weight compared to WT fish (Fig. 2A). Locomotor activity of adult fish was unchanged (Fig. S1C). Similar to mouse *Tmbim5*, zebrafish *tmbim5* is expressed in most tissues with the highest mRNA expression levels in brain and muscle (Fig. 2B). A histopathological examination of these tissues and liver revealed no major differences (Fig. S2A-J).

**Figure 2.**
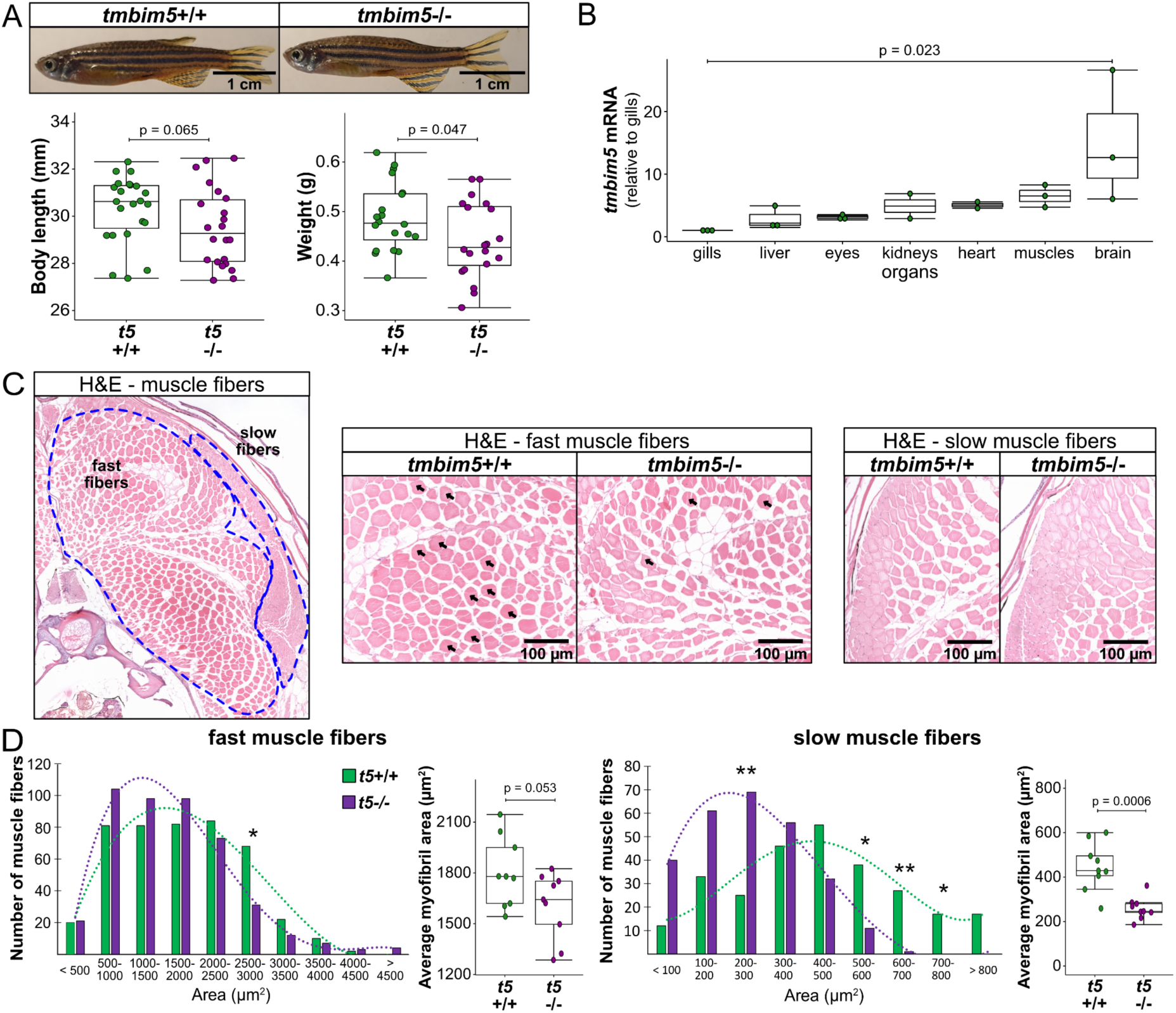
Tmbim5 deficiency causes muscle atrophy with preferential effects on slow-twitch fibers. A) Smaller size of *tmbim5*^-/-^ adult fish. Representative images of WT and *tmbim5*^-/-^ 8-month-old zebrafish. Morphometric analysis results are presented as box-and-whisker plots (box: 25th–75th percentile; whiskers: min to max), with each dot representing an individual fish (*n* = 19–23). Number of experiments = 3 (weight) or 4 (body length). Statistical analysis: Mann–Whitney test (body length) or *t*-test (weight). B) *tmbim5* mRNA levels in various adult zebrafish organs measured using qPCR. Expression was normalized to the gills (the organ with the lowest *tmbim5* expression), with *rpl13a* used as a reference gene. Data are presented as box-and-whisker plots, with each dot representing an independent biological replicate (*n* = 2–3; each RNA sample was isolated from 2 fish). Number of experiments = 3. Statistical analysis: Kruskal-Wallis test followed by Dunn’s test. C) Representative images of hematoxylin and eosin (H&E)-stained skeletal muscle sections from adult (8-month-old) zebrafish with fast- and slow-twitch fibers indicated. Middle panel: magnification of the area with fast fibers images from WT and *tmbim5*^-/-^. Arrows indicate fibers of 2500-3000 µm², which are significantly reduced in number in the mutants. Right panel: magnification of the area with slow-twitch fibers images of WT and *tmbim5*^-/-^. D) Smaller cross-sectional area of fast and slow muscle fibers in *tmbim5*^-/-^ adult fish. Histograms showing the distribution of fast (left panel) and slow (right panel) muscle fiber cross-sectional areas in WT and *tmbim5*^-/-^ fish. The mutant distribution is shifted toward smaller fibers. Statistical analysis: Chi-square test (fast fibers: *p* = 0.0009; slow fibers: *p* < 0.0001). Differences in the number of fibers of specific sizes are marked with an asterisk (* *p* < 0.05), analyzed using the Mann–Whitney test or *t*-test with Benjamini-Hochberg (BH) correction for multiple comparisons. Box-and-whisker plots showing the average myofibril area. Each dot represents the average for one section (50 fibers analyzed per section, *n* = 9). Number of fish = 3. Statistical analysis: *t*-test.

Because of the muscle phenotype implied by reduced tail coiling and delayed hatching of larvae, we next analyzed the skeletal muscle in more detail. In zebrafish slow (red) and fast (white) twitch fibers are segregated, allowing analysis without immunohistochemical staining ^35^ (Fig. 2C). Slow twitch fibers are rich in mitochondria and depend more on aerobic metabolism than fast (white) twitch muscle fibers ^36^. A morphometric analysis of these fibers demonstrated a shift towards smaller fibers, a decline in the fibers with a cross-sectional area of 2500-3000 µm^2^, and a reduction in the average cross-sectional area in mutant fish. All of these changes were more pronounced in slow twitch muscle fibers than in fast twitch fibers (Fig. 2D). The more pronounced atrophy of slow-twitch muscle fibers possibly reflects mitochondrial dysfunction and is consistent with the reduced muscle strength observed in larvae.

### Tmbim5 does not function as an independent mitochondrial Ca²⁺ uptake pathway

The MCU complex is the major mitochondrial Ca²⁺ uptake mechanism ^1,2^. To clarify whether Tmbim5, as overexpression results in human embryonic kidney cells suggested ^20^, can act as an additional Ca^2+^ uptake mechanism besides the MCU complex or modify MCU activity, we crossed *tmbim5*^−/−^ mutants with *mcu* KO fish. We hypothesized that depletion of two potentially independent uptake mechanisms would be detrimental and/or alter mitochondrial Ca^2+^ uptake. *mcu* KO fish lack the capacity to uptake mitochondrial Ca^2+^ but have an otherwise unremarkable phenotype ^11^.

We found no differences in the distribution of adult zebrafish genotypes following a heterozygous cross (Fig. 3A) suggesting no additive detrimental effect on development. We also measured mitochondrial Ca^2+^ uptake in isolated mitochondria by quantifying external Ca^2+^ levels in the bath (Fig. 3B) These experiments confirmed the previous observation that mitochondria from Mcu-deficient zebrafish show almost no Ca^2+^ uptake ^11^. There was no additional effect of *tmbim5* knockout on Ca^2+^ uptake. Together these results imply that Tmbim5 does not constitute a relevant mitochondrial Ca^2+^ uptake system independent of Mcu. It does not rule out a modulatory effect of Tmbim5 on the MCU complex.

**Figure 3.**
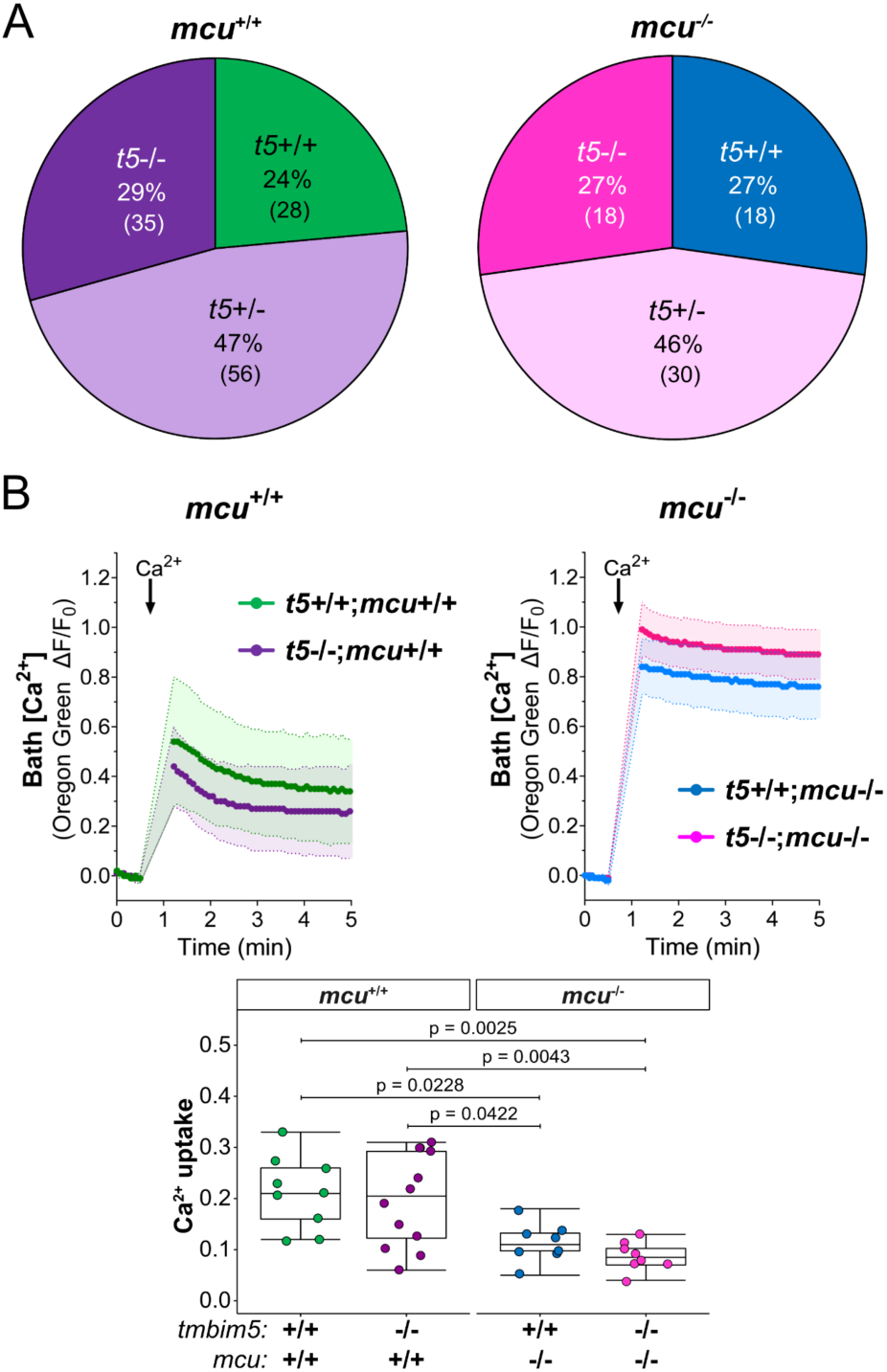
Tmbim5 does not function as an independent mitochondrial Ca²⁺ uptake pathway. A) Genotype distribution does not deviate from the expected Mendelian ratio after *tmbim5*^+/-^*;mcu*^+/-^ zebrafish breeding. *n* is indicated in brackets. Statistical analysis: Chi-square test. B) Loss of Tmbim5 does not significantly affect mitochondrial Ca²⁺ uptake in mitochondria isolated from *mcu*^-/-^ larvae. Mitochondria were loaded with Ca^2+^ (20 µM), revealing significantly reduced Ca^2+^ uptake in both single *mcu*^-/-^ and double *tmbim5*^-/-^*;mcu*^-/-^ knockouts. Top: fluorescence of Oregon Green 488 BAPTA-5N was normalized to the initial value (F_0_). Data are presented as mean ± SD ΔF/F_0_ traces. Gaps in the measurements are due to technical limitations of the equipment and procedure, which required pauses in fluorescence readings during reagent addition. Bottom: quantification of Ca^2+^ uptake, calculated within 2 min after CaCl_2_ addition as the difference between the maximum ΔF/F_0_ value after Ca^2+^ addition and the ΔF/F_0_ value reached 2 min later. Results are presented as box-and-whisker plots (box: 25th–75th percentile; whiskers: min to max), with each dot representing an independent biological replicate (*n* = 8–12). Statistical analysis: one-way ANOVA followed by post-hoc Tukey HSD test.

### Zebrafish Slc8b1 is the likely ortholog of the mitochondrial Na⁺/Ca²⁺ exchanger NCLX

TMBIM5 might also constitute an additional mitochondrial Ca^2+^ efflux system ^28,29^ besides sodium-dependent mitochondrial Ca^2+^ release mediated by NCLX ^17^. To clarify whether double knockout of both proteins would be detrimental, we first had to generate NCLX-deficient fish. Unfortunately, the identity of NCLX in fish is unknown. However, the *slc8b1* gene encodes a protein of 308 amino acids with the highest homology (24%) to human NCLX and a predicted sodium/calcium exchanger domain at its N-terminus (Fig. S3A). It is noteworthy that the predicted fish protein is only half as big as human NCLX which has a length of 584 amino acids.

We designed a gRNA to target the conserved exchanger domain which led to the insertion of 17 bp resulting in a frameshift mutation, disrupting the proposed transporter function of Slc8b1 (Fig. S3B). This resulted in a decrease in *slc8b1* mRNA levels. There were no compensatory changes of *tmbim5* mRNA (Fig. S3C). *slc8b1* (*nclx*) mRNA levels peaked at six hours post fertilization and declined thereafter in wildtype embryos (Fig. 4A). *In situ* hybridization revealed that *slc8b1* is strongly expressed in the brain and in neuromasts, which are sensory patches distributed on the head and along the lateral line of fish ^37^ (Fig. 4B). In adult fish, *tmbim5* and *slc8b1* are both highly expressed in the brain but *slc8b1* is expressed at a low level in skeletal muscles (Fig. 4C). There is therefore some but no complete overlap of the expression patterns of *tmbim5* and *slc8b1*.

**Figure 4.**
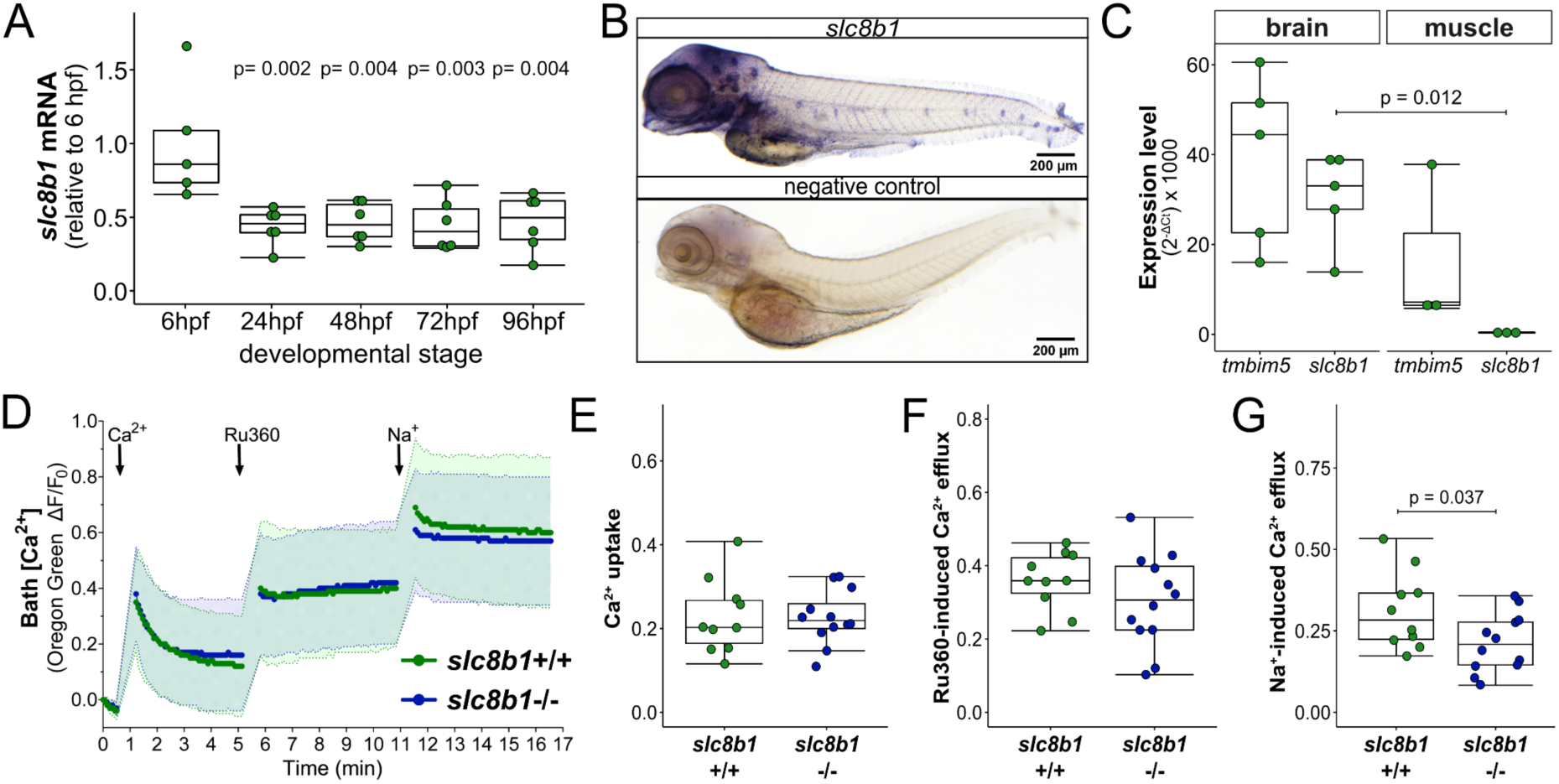
Zebrafish Slc8b1 is the likely ortholog of the mitochondrial Na⁺/Ca²⁺ exchanger NCLX. A) *slc8b1* mRNA expression at different developmental stages, quantified using qPCR. Expression was normalized to 6 hpf, with *18S* used as a reference gene. Results are shown as box-and-whisker plots (box: 25th–75th percentile, whiskers: min–max), with each dot representing an independent biological replicate (*n* = 5–6, each RNA sample was isolated from 30 embryos/larvae). Statistical analysis: one-way ANOVA with post-hoc Tukey HSD test (*p*-values for comparisons with 6 hpf). B) *slc8b1* transcript detected by whole-mount in situ hybridization in 4 dpf wild-type zebrafish. A sense probe was used as a negative control. Number of experiments = 3. C) Comparative expression of *slc8b1* and *tmbim5* in the brain and skeletal muscle of adult zebrafish, assessed by qPCR. Data are presented as box-and-whisker plots, each dot representing an independent biological replicate (*n* = 3–5, each RNA sample pooled from 2 fish). Statistical analysis: *t*-test with Benjamini-Hochberg (BH) correction for multiple comparisons. D) Functional analysis of mitochondrial Ca^2+^ transport in mitochondria isolated from *slc8b1*^-/-^ and WT larvae, using Oregon Green 488 BAPTA-5N fluorescence measurements Changes in bath Ca^2+^ levels measured under basal conditions and after treatment with Ru360 (10 µM) to inhibit mitochondrial Ca^2+^ uptake. Upon addition of Na^+^ (10 mM), a reduced Na^+^-dependent Ca^2+^ efflux was observed in *slc8b1*^-/-^ mitochondria. Fluorescence was normalized to initial values (F_0_), and data are presented as mean ± SD ΔF/F_0_ traces. E) *slc8b1* knockout does not affect Ca^2+^ uptake into mitochondria. Quantification of mitochondrial Ca^2+^ uptake was performed 2 min after CaCl_2_ addition. F) No significant difference in Ca^2+^ efflux following Ru360 treatment. Efflux was estimated by subtracting the last ΔF/F_0_ value after CaCl_2_ addition from the maximum ΔF/F_0_ after Ru360 addition. G) Reduced Ca²⁺ efflux from *slc8b1^-/-^* mitochondria after Na⁺ addition compared to WT. Efflux was quantified as the difference between the last ΔF/F_0_ value after Ru360 addition and the maximum ΔF/F_0_ after Na^+^ addition. E-G) Results are presented as box-and-whisker plots, with each dot representing an independent biological replicate (*n* = 10–12). Statistical analysis: *t*-test.

To assess whether Slc8b1 mediates Na⁺-dependent Ca^2+^ efflux from mitochondria, we measured bath Ca^2+^ levels in mitochondria isolated from WT and *slc8b1*^−/−^ larvae. After loading mitochondria with Ca^2+^, MCU-mediated uptake was inhibited with Ru360, followed by stimulation of Na^+^-dependent Ca^2+^ efflux with a Na^+^-containing buffer (Fig. 4D). *slc8b1*^−/−^ mitochondria had a similar Ca^2+^ uptake (Fig. 4E) and response to Ru360 (Fig. 4F) to WT. However, sodium-dependent Ca^2+^ efflux (Fig. 4G) was significantly but not completely impaired supporting Slc8b1 as a key component of this process and a likely NCLX ortholog.

### Combined loss of Tmbim5 and Slc8b1 severely disrupts mitochondrial Ca^2+^ homeostasis

Having established Slc8b1 as a relevant mitochondrial Ca^2+^ efflux system, we next examined whether removing both transport systems would be deleterious. The Mendelian distribution of offspring was as expected (Fig. 5A), indicating that the double knockout was not embryonically lethal. However, quantitative assessment of Ca^2+^ fluxes revealed significant perturbations in *tmbim5*/*slc8b1* double knockout mitochondria obtained from larvae (Fig. 5B).

**Figure 5.**
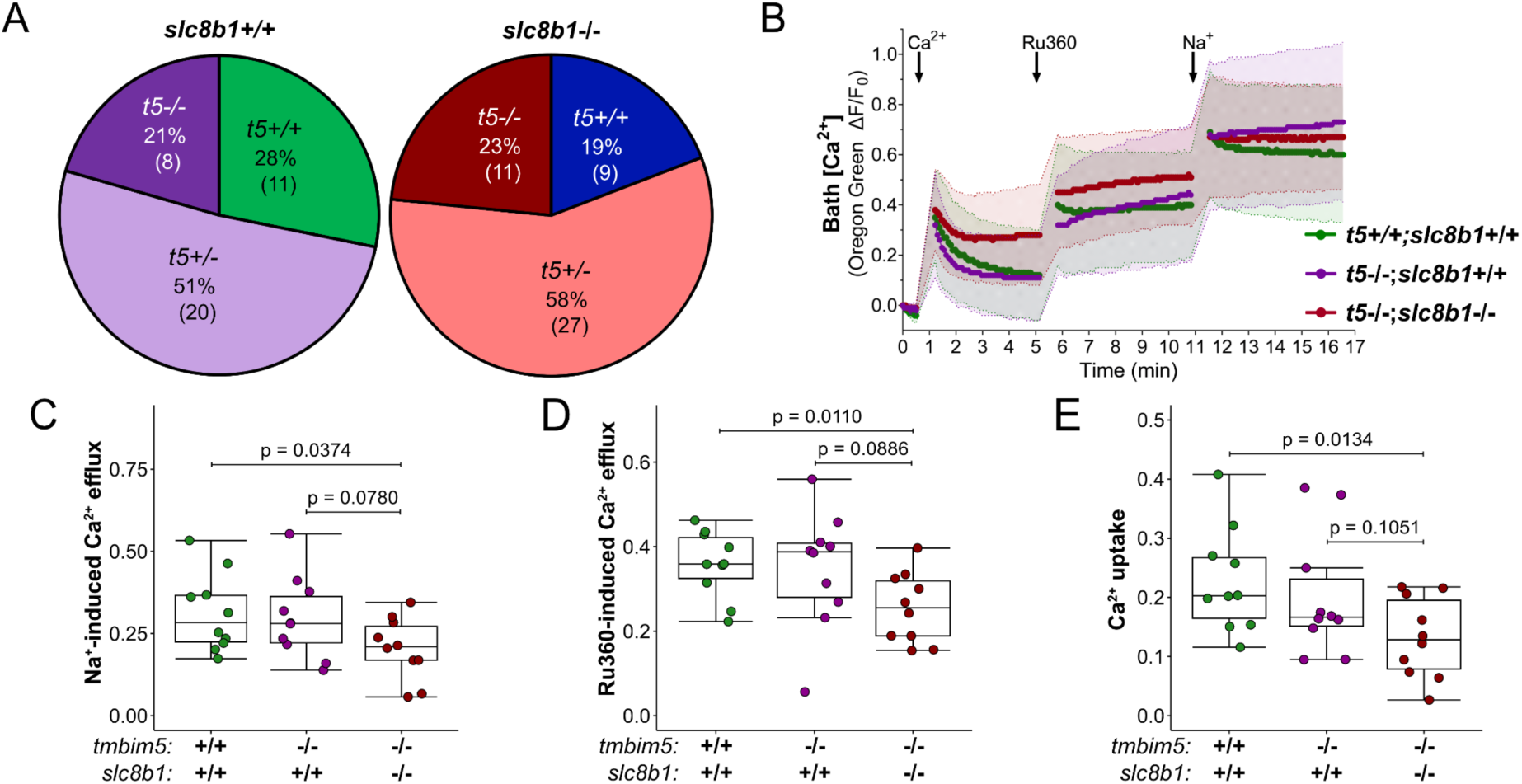
Combined loss of Tmbim5 and Slc8b1 severely disrupts mitochondrial Ca^2+^ homeostasis. A) Genotype distribution of offspring from heterozygous *tmbim5*^+/-^*;slc8b1*^+/-^ zebrafish breeding does not differ from expected Mendelian ratios. Sample sizes are indicated in brackets. Statistical analysis: Chi-square test, number of experiments = 3. B) Changes in bath Ca^2+^ measured by Oregon Green 488 BAPTA-5N in mitochondria isolated from wild-type (WT), *tmbim5*^-/-^, and *tmbim5*^-/-^*;slc8b1*^-/-^ larvae. Mitochondria were first loaded with Ca^2+^ (20 µM), followed by the addition of Ru360 (10 µM) to inhibit Ca^2+^ influx. Next, Na^+^ (10 mM) was added to assess Ca^2+^ efflux via NCLX activity, showing reduced Na^+^-dependent Ca^2+^ efflux in *slc8b1^-/-^*mitochondria. Fluorescence was normalized to the initial value (F_0_). Data are presented as mean ± SD ΔF/F_0_ traces. C) Reduced Na^+^-dependent Ca^2+^ efflux in mitochondria from *tmbim5*^-/-^*;slc8b1*^-/-^ larvae compared to WT. The amount of Ca^2+^ efflux following Na^+^ addition was estimated by subtracting the last ΔF/F_0_ value obtained after Ru360 addition from the maximal ΔF/F_0_ value after Na^+^ addition. D) Decreased Ca^2+^ efflux from mitochondria isolated from *tmbim5^-/-^;slc8b1^-/-^*larvae. The amount of Ca^2+^ efflux after Ru360 addition was estimated by subtracting the last ΔF/F_0_ value obtained after CaCl_2_ addition from the maximal ΔF/F_0_ value after Ru360 addition. E) Reduced Ca^2+^ uptake by mitochondria from *tmbim5*^-/-^*;slc8b1*^-/-^ larvae compared to WT. Ca^2+^ uptake was quantified 2 minutes after CaCl_2_ addition. C-E) Results are presented as box-and-whisker plots (box: 25th–75th percentile, whiskers: min–max), with each dot representing an independent biological replicate (*n* = 10). Statistical analysis: two-way ANOVA followed by *t*-test or Mann–Whitney test.

Consistent with findings from mammalian cells, where combined TMBIM5 and NCLX dysfunction compromised mitochondrial Ca^2+^ handling capacity ^28,29^ and impaired ruthenium red-sensitive Ca^2+^ efflux ^28^, we observed similar alterations. Both Na^+^-stimulated efflux (Fig. 5C) and efflux after blocking further import with Ru360 (Fig. 5D) were reduced. Additionally, the capacity for Ca^2+^ uptake was decreased in *tmbim5*/*slc8b1* double knockout mitochondria (Fig. 5E), possibly due to a reduced electrochemical driving force. These findings from purified mitochondria obtained from whole larvae support the conclusion that Tmbim5 functions as an additional mitochondrial Ca^2+^ efflux system alongside Slc8b1. With this functional validation established, we proceeded to examine the organismal-level phenotypic consequences of the double knockout.

### Tissue-specific effects of dual Ca^2+^ transport system disruption

We investigated the effect of double knockout on the phenotypes associated with Tmbim5 loss of function to understand how disruption of multiple Ca^2+^ transport pathways affects different tissues. Similarly to *tmbim5*^−/−^ larvae (Fig. 1E), *slc8b1*^−/−^ and *tmbim5*/*slc8b1* KO larvae exhibited a lower hatching efficiency compared to WT (Fig. 6A). Also, coiling activity, which precedes hatching, was reduced in *tmbim5*/*slc8b1* embryos compared to wildtype and *slc8b1*^−/−^ (Fig. 6B) suggesting skeletal muscle impairment. This correlated with pronounced alterations in mitochondrial ultrastructure in double knockout animals shown by transmission electron microscopy (Fig. 6C). Despite these changes, spontaneous (Fig. S4A) and light-induced (Fig. S4B) locomotor activity of *slc8b1*^−/−^ and *tmbim5*/*slc8b1* larvae were normal.

**Figure 6.**
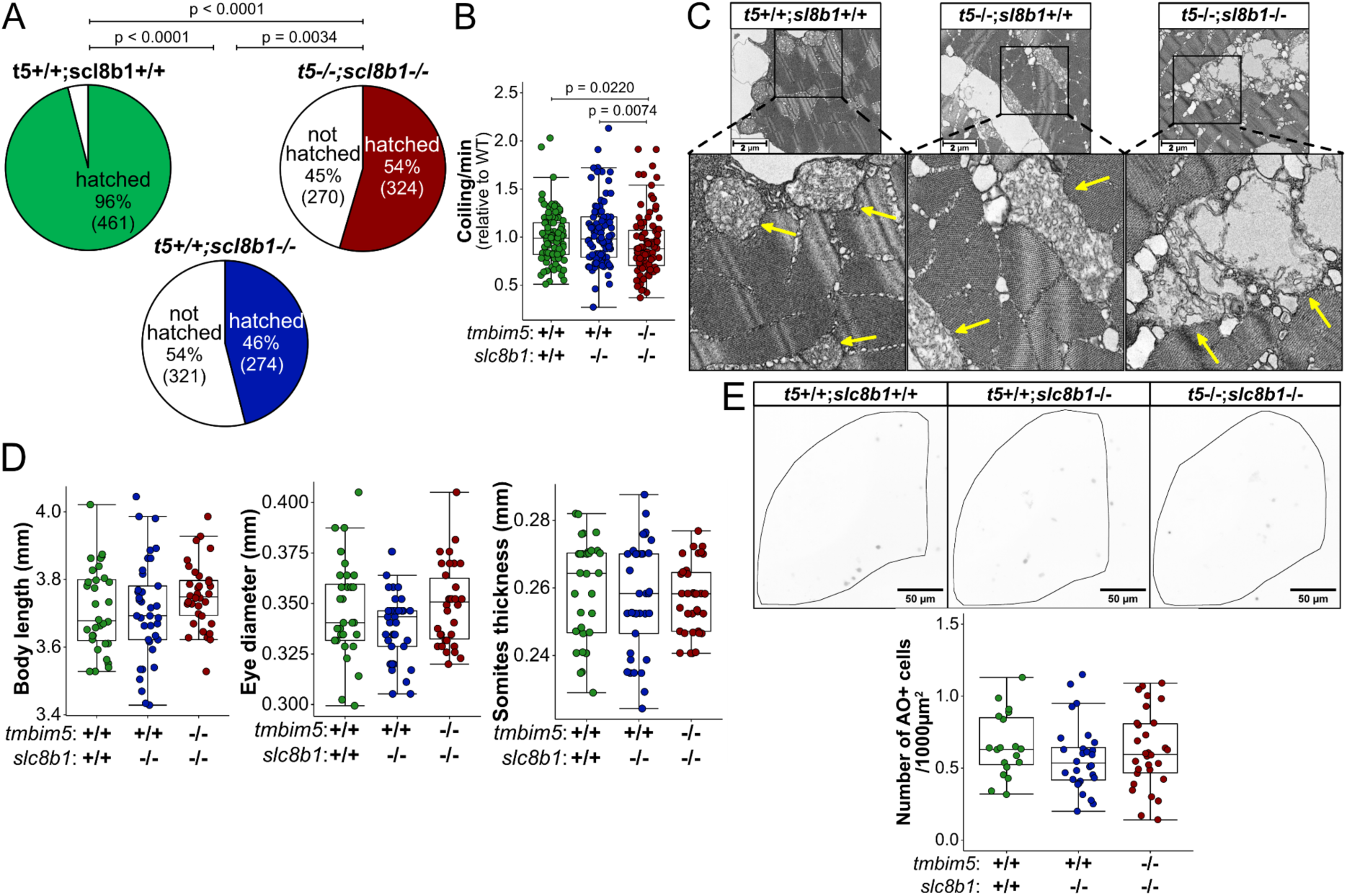
Tissue-specific effects of dual Ca^2+^ transport system disruption. A) Reduced hatching efficiency of *slc8b1*^-/-^ and *tmbim5*^-/-^*;slc8b1*^-/-^ larvae. *n* is indicated in brackets. Number of experiments = 3. Statistical analysis: Chi-square test. B) Decreased coiling activity of *tmbim5*^-/-^*;slc8b1*^-/-^ embryos at 1 dpf. The mean number of coiling movements per minute, normalized to WT from the same experiment, is shown as box-and-whisker plots (box: 25th–75th percentile; whiskers: min to max). Each dot represents an individual embryo (*n* = 94–96). Number of experiments = 3. Statistical analysis: two-way ANOVA followed by Mann–Whitney test. C) Aberrant mitochondrial morphology in *tmbim5^-/-^;slc8b1^-/-^* in the muscle of double knockout larvae. Representative transmission electron microscopy images of mitochondria from skeletal muscle of 5 dpf WT, *tmbim5^-/-^*, and *tmbim5^-/-^;slc8b1^-/-^* zebrafish larvae. Arrows indicate mitochondria in magnified images. D) Normal morphology and body size of *slc8b1*^-/-^ and *tmbim5*^-/-^*;slc8b1*^-/-^ larvae at 5 dpf. Morphometric measurements (body length, eye diameter, and somite thickness) are presented as box-and-whisker plots, with each dot representing an individual larva (*n* = 32–34). Statistical analysis: one-way ANOVA with post-hoc Tukey HSD test (body length and eye diameter) or Kruskal-Wallis test with Dunn’s post-hoc test (somite thickness). Number of experiments = 3. E) No increased cell death in the optic tectum of *slc8b1*^-/-^ or *tmbim5^-/-^;slc8b1*^-/-^ larvae analyzed with AO staining. Representative images of WT and *slc8b1*^-/-^ and *tmbim5*^-/-^*;slc8b1*^-/-^ larvae (black dots indicate high AO signals). Quantification results are presented as box-and-whisker plots, with each dot representing an individual larva (*n* = 19–30, number of experiments = 3). Statistical analysis: two-way ANOVA.

Remarkably, all phenotypes associated with reduced size i.e. larval body length, eye diameter and somite thickness were surprisingly similar between wildtype, *slc8b1*^−/−^ and *tmbim5*/*slc8b1* double knockout larvae (Fig. 6D). Double knockout also rescued the increased cell death in the central nervous system observed in *tmbim5*^−/−^ larvae (Fig. 6E). Together these data imply that concurrent loss of both transport systems affects different tissues differentially. While the effects on coiling were more pronounced, corresponding to severe disruption of the mitochondrial cristae architecture, the effects of *tmbim5* loss of function on body size and increased cell death in nervous tissue were rescued by concomitant knockout of *tmbim5* and *slc8b1*. This suggests tissue-specificity of mitochondrial Ca^2+^ transport systems or tissue-specific additional function of Tmbim5 or Slc8b1.

### Mitochondrial Ca^2+^ and membrane potential changes reveal tissue-specific compensation mechanisms

To address this tissue specificity, we therefore decided to assess changes in steady-state mitochondrial Ca^2+^ and membrane potential levels of WT, *slc8b1*^−/−^ and *tmbim5*/*slc8b1* KO larvae in susceptible and non-susceptible tissues, here brain and muscle. To quantify basal mitochondrial Ca^2+^ levels, we injected plasmid DNA encoding the 4mtD3cpv Ca^2+^ probe into zebrafish embryos ^38^. This FRET-based sensor allows ratiometric imaging and quantification of Ca^2+^ amounts independent of probe expression level ^39,40^. To quantify mitochondrial membrane potential, we injected tetramethylrhodamine ethyl ester (TMRE) into the yolk of 1-cell stage zebrafish embryos as described in Vicente et al. ^41^. TMRE is a positively charged dye known to accumulate in mitochondria due to their membrane potential. Two days after injection, we recorded a TMRE signal in brain and skeletal muscle that was significantly reduced after depolarization with carbonyl cyanide 3-chlorophenylhydrazone (CCCP) used as positive control.

In the brain, we did not observe any significant changes in basal mitochondrial Ca^2+^ levels of *tmbim5*^−/−^ or *tmbim5*/*slc8b1* double knockout larvae (Fig. 7A). However, mitochondrial membrane potential was reduced in Tmbim5-deficient larvae and rescued in *tmbim5*/*slc8b1* double knockout (Fig. 7B). In the muscles of *tmbim5*/*slc8b1* larvae, in contrast, the mitochondrial Ca^2+^ content was significantly lower than in WT or *tmbim5* single knockout (Fig. 7C) in line with the reduced mitochondrial Ca^2+^ uptake observed in purified mitochondria (Fig. 5E). MMP was similar between the genotypes (Fig. 7D) ruling out changes in mitochondrial Ca^2+^ uptake mediated by changes of the driving force. These results therefore corroborate our phenotypic analysis where double knockout of *tmbim5* and *slc8b1* aggravates or ameliorates Tmbim5 loss of function in a tissue-specific manner. The beneficial effect in the brain appears not to be mediated by changes in mitochondrial Ca^2+^ homeostasis but rather via effects on the activity of the electron transfer system possibly by changes of AFG3L2-mediated remodeling of the respiratory complexes. In muscle, the opposite appears to be true.

**Figure 7.**
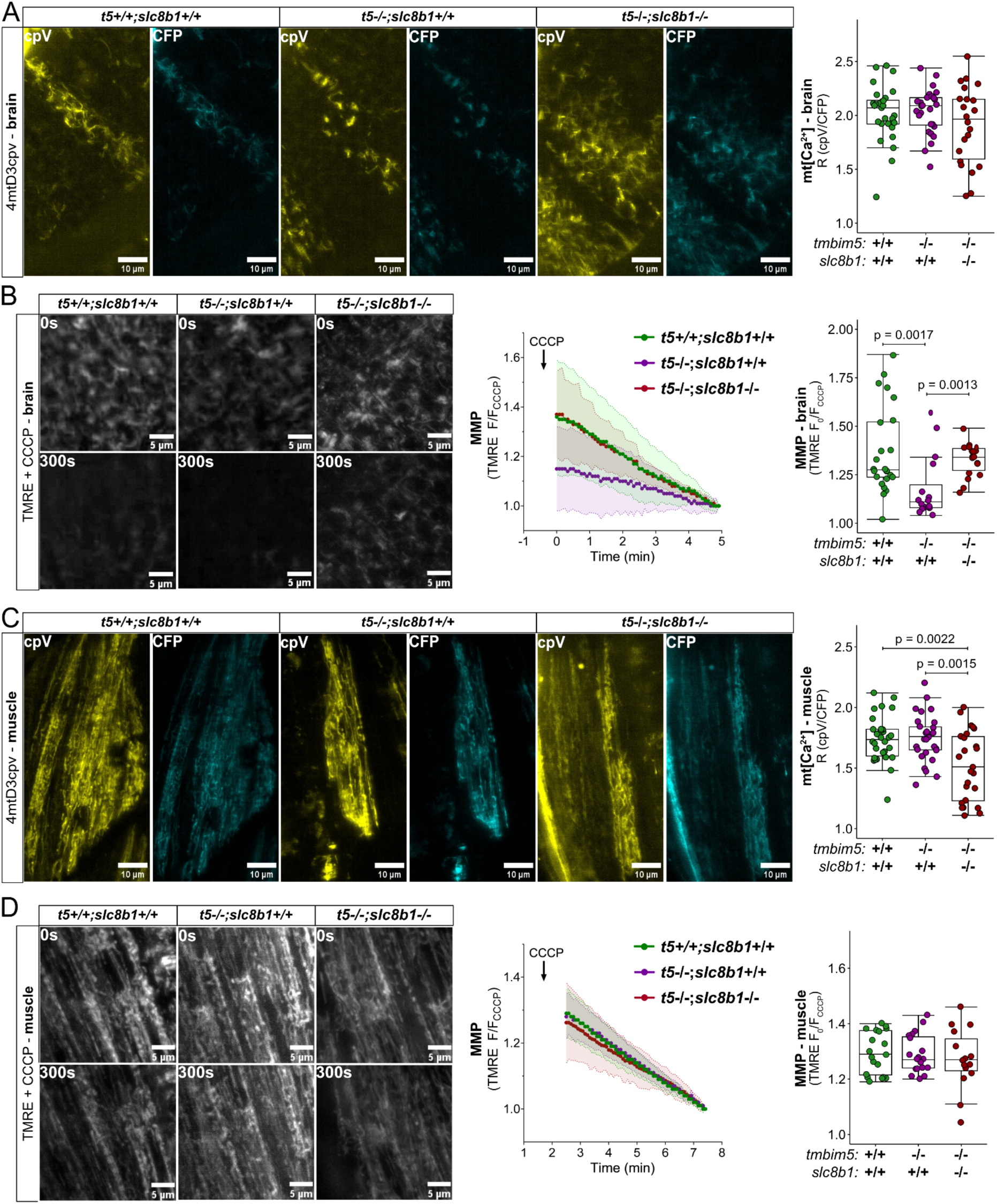
Mitochondrial Ca²⁺ and membrane potential changes reveal tissue-specific compensation mechanisms. A) Basal mitochondrial Ca^2+^ level in the brain is not affected by knockout of *tmbim5* or *tmbim5*/*slc8b1* double knockout larvae. Left: representative images of brain (optic tectum) of 2 dpf larvae with transient and mosaic expression of the 4mtD3cpv Ca^2+^ probe. Right: mitochondrial [Ca^2+^], assessed from the ratio of cpV to CFP (R), is shown as a box-and-whisker plot (box: 25th-75th percentile; whiskers: min to max), with each point representing an individual larva (*n* = 22-29). Statistical analysis: two-way ANOVA followed by *t*-test. B) Decreased mitochondrial membrane potential in *tmbim5*^-/-^, but not in *tmbim5*^-/-^*;slc8b1*^-/-^ zebrafish observed *in vivo* in the optic tectum of 2 dpf larvae. Zebrafish embryos were injected with TMRE (1 nl of 2 mM) at the 1-cell stage and imaged *in vivo* using LSFM at 2 dpf. CCCP (10 µM) was used to depolarize mitochondria, and the F_0_/F_CCCP_ ratio was calculated. Left: representative images of the ROI in the brain at the beginning and after 5 min of CCCP treatment. Right: changes in membrane potential (MMP) measured in the brain by TMRE. Data are presented as mean ± SD F/F_CCCP_. MMP, calculated as the F_0_/F_CCCP_ ratio, is plotted as box-and-whisker plots, with each dot representing an individual larva (*n* = 16–24). Statistical analysis: two-way ANOVA followed by Mann–Whitney test. C) Decreased basal mitochondrial Ca^2+^ level in muscles of *tmbim5*^-/-^;*slc8b1*^-/-^ larvae. Left: representative images of muscle of 2 dpf larvae with transient and mosaic expression of the 4mtD3cpv Ca^2+^ probe. Right: mitochondrial [Ca^2+^], assessed as in A), is shown as a box-and-whisker plot, with each point representing an individual larva (n = 25-30). Statistical analysis: two-way ANOVA followed by t-test. D) Unaffected mitochondrial membrane potential in *tmbim5*^-/-^ and in *tmbim5*^-/-^;*slc8b1*^-/-^ zebrafish measured *in vivo* in the muscle of 2 dpf larvae. Measurements were performed as described in B). Left: representative images of the ROI in the muscle at the beginning and after 5 min of CCCP treatment. Right: changes in MMP measured in the muscle by TMRE. Data are presented as mean ± SD F/F_CCCP_. MMP, calculated as the F_0_/F_CCCP_ ratio, is plotted as box-and-whisker plots, with each dot representing an individual larva (*n* = 15–20). Statistical analysis: two-way ANOVA followed by Mann–Whitney test.

Our results are therefore consistent with TMBIM5 functioning as a mitochondrial Ca^2+^ efflux pathway, as evidenced by the impaired Ca^2+^ handling capacity when both Tmbim5 and Slc8b1 are absent. The dysregulated mitochondrial Ca^2+^ homeostasis resulting from simultaneous disruption of these efflux pathways may directly contribute to the observed ultrastructural abnormalities. The disconnect between severe subcellular abnormalities and mild organismal phenotypes underscores the remarkable adaptability of mitochondrial Ca^2+^ homeostatic systems *in vivo*.

### Transcriptional changes of genes encoding the mitochondrial Ca^2+^ machinery differ between brain and skeletal muscle

To investigate whether the differential effects of Tmbim5 and Slc8b1 loss in brain and muscle result from compensatory transcriptional changes, we analyzed the expression levels of genes involved in mitochondrial Ca^2+^ handling: *mcu*, *micu1–3*, *efhd1*, *letm1*, *letm2*, *letmd1*, *slc8b1*, *tmem65*, and *afg3l2*. *slc25a28*, encoding a Ca^2+^-independent iron transporter served as control. We compared expression between wildtype and *tmbim5* KO, wildtype and *tmbim5/slc8b1* double knockout, and *tmbim5* KO versus double knockout in both tissues.

IIn the brain, *micu3* was significantly downregulated in *tmbim5* KO (Fig. 8A). Double knockout (*tmbim5/slc8b1*) resulted in downregulated *efhd1* and *letm2* levels compared to wild-type, while *tmem65* was specifically reduced compared to *tmbim5* KO (Fig. 8A). In skeletal muscle, *tmbim5* KO reduced *mcu* and *letm2* while increasing *efhd1* (Fig. 8B). Conversely, double knockout showed *letm2* upregulation compared to both wild-type and *tmbim5* KO (Fig. 8B). Notably, the expression levels of all mitochondrial calcium machinery genes were broadly reduced in double knockout brain tissue (two-way ANOVA: *p*=0.000002, Fig. 8C) and in *tmbim5* KO muscle, although less evidently (*p*=0.01816, Fig. 8D). These changes corresponded to a significant reduction in mitochondrial DNA content in the brains of Tmbim5-depleted fish that persisted in the *tmbim5*/*slc8b1* double KO (Fig. 8E) but not in muscle (Fig. 8F).

**Figure 8.**
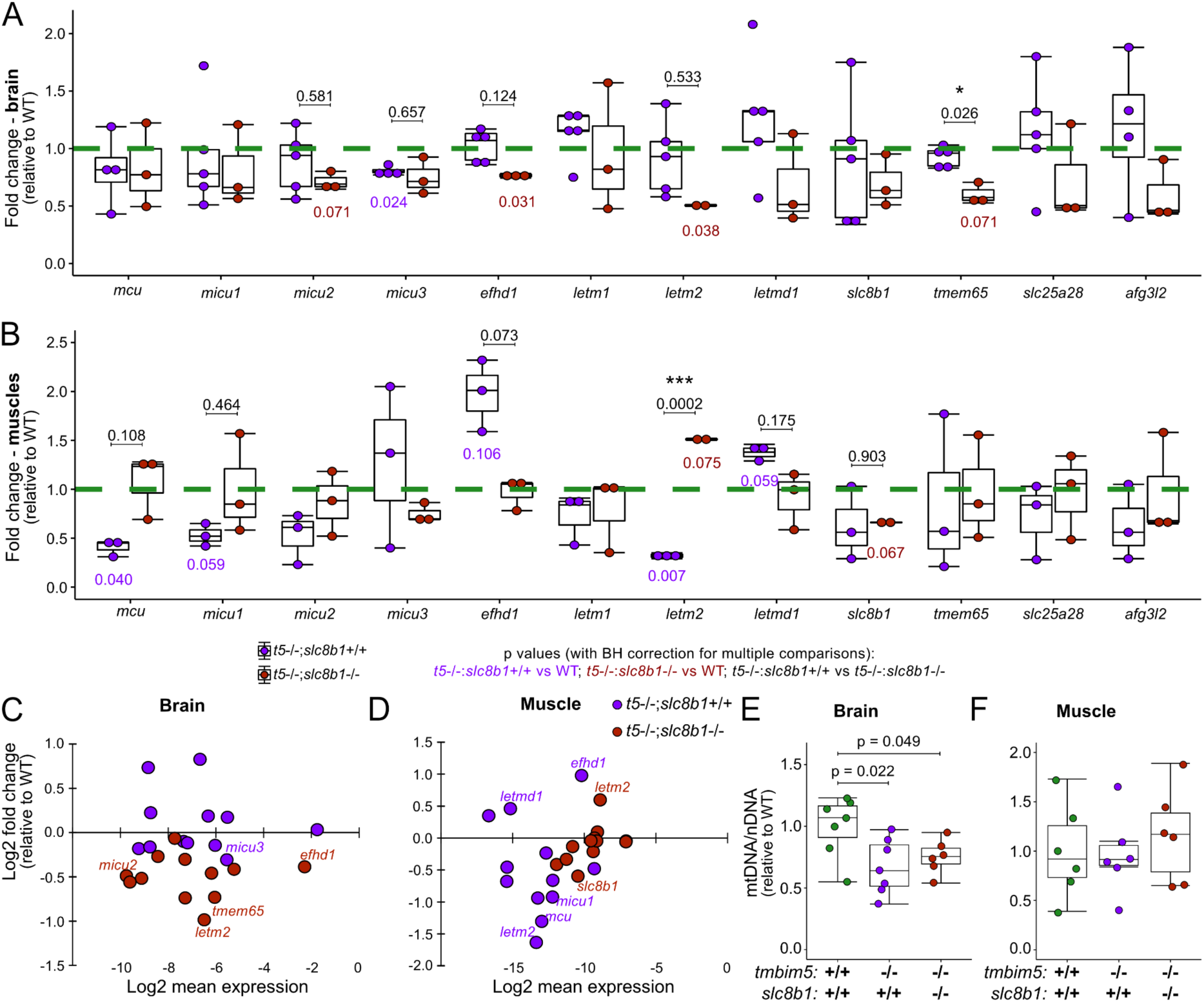
Transcriptional changes of genes encoding the mitochondrial Ca^2+^ machinery differ between brain and skeletal muscle. A) *micu3* is downregulated in the brain of *tmbim5*^-/-^ fish, while *efhd1* and *letm2* expression is reduced in the brain of *tmbim5*^-/-^;*slc8b1*^-/-^. B) In the muscle of *tmbim5*^-/-^ fish, *mcu* and *letm2* are downregulated; however, in *tmbim5*^-/-^;*slc8b1*^-/-^ fish, *letm2* is upregulated in the muscle. C-D) qPCR results presented as MA plots showing the relation between fold change and the mean expression levels of examined genes in the brain (C) and muscle (D) calculated as 2^(-ΔCq)^. A-D) mRNA levels of proteins involved in mitochondrial Ca^2+^ transport were quantified by qPCR in the brain and skeletal muscles of adult (8-14-month-old) zebrafish and normalized to WT. *rpl13a* and *ef1a* were used as reference genes. Results are presented as box-and-whisker plots (box: 25th–75th percentile; whiskers: min to max), with each dot representing an independent biological replicate (*n* = 2–6, with each RNA sample isolated from 2 fish). Number of experiments = 3-6. Statistical analysis: one-sample *t*-test (comparisons with WT) or *t*-test with BH correction for multiple comparisons. *p*-values shown below the boxplots indicate comparisons with WT: magenta values represent comparisons with *tmbim5* knockout, and red values represent comparisons with *tmbim5/slc8b1* double knockout. Black *p*-values above boxplots indicate comparisons between *tmbim5* knockout and *tmbim5/slc8b1* double knockout. E) The mitochondrial (mtDNA) to nuclear DNA (nDNA) ratio is decreased in the brain of *tmbim5*^-/-^ and *tmbim5*^-/-^;*slc8b1*^-/-^ zebrafish. F) Unchanged mtDNA to nDNA ratio in the skeletal muscle of *tmbim5*^-/-^ and *tmbim5*^-/-^;*slc8b1*^-/-^ zebrafish. E-F) The mtDNA/nDNA ratio was quantified by qPCR and normalized to WT. *mt-nd1* (*mitochondrial NADH dehydrogenase 1*) was used as a marker for mtDNA, while *ef1a* was used for nDNA. Results are presented as box-and-whisker plots, with each dot representing an independent biological replicate (*n* = 6). Number of experiments = 2. Statistical analysis: two-way ANOVA followed by *t* test.

The downregulation of *micu3* in brain and *mcu* in muscle as well as the upregulation of *efhd1*, which is an inhibitor of the MCU ^43^, is in line with compensatory changes reducing mitochondrial Ca^2+^ influx suggesting that Tmbim5 is mainly part of an efflux system. We posit that *letm2* may mediate tissue-specific effects, being downregulated in protective brain but upregulated in affected muscle. While previously being reported as testis-specific ^44^, *letm2* is expressed in the brain (Fig. 8B) and even dominates over *letm1* in zebrafish muscle (Fig. 8D). Not much is known about its function. Alternatively, coordinated downregulation of the entire mitochondrial Ca^2+^ machinery may protect brain tissue against *tmbim5* deficiency by shifting energy supply to glycolysis as described for astrocyte-specific *Nclx* knockout in mice ^45^.

## Discussion

Our findings establish Tmbim5 as critical for growth and muscle function in zebrafish and demonstrate that it functions predominantly as a mitochondrial Ca^2+^ efflux rather than influx transport system. This conclusion is based on the lack of additive effects in *tmbim5*/*mcu* double knockouts and the functional impairment when both efflux pathways (*tmbim5* and Na^+^-dependent Ca^2+^ efflux via *slc8b1*) were disrupted simultaneously. Surprisingly, *slc8b1* knockout alone produced minimal phenotypic effects, contrasting with the embryonic lethality observed in mammalian *NCLX* knockouts. Even more unexpected, the double knockout of both efflux pathways was protective in some tissues but detrimental in others. These results suggest that the functions of Tmbim5 and Slc8b1 can be tissue-specific and indicate the existence of additional, yet uncharacterized mitochondrial Ca^2+^ transport mechanisms.

Tmbim5 deficiency decreased the size of zebrafish larvae and weight of adults in line with knockdown of the *tmbim5* ortholog in snails ^46^ but not in mice ^27^. However, similar to *Tmbim5* mutant mice that suffer from myopathy, myofiber cross-sections were smaller in *tmbim5*^-/-^ fish, indicating muscle atrophy. This was particularly pronounced in slow-twitch muscles, which contain abundant mitochondria and rely heavily on oxidative phosphorylation compared to fast-twitch fibers ^36^, suggesting a link with compromised mitochondrial function. These pathological alterations of muscle function probably underlie the delayed escape of Tmbim5-deficient embryos from the chorion which is mediated by tail coiling ^34^. Interestingly, *Mcu* knockout in mice also negatively affects size ^8^ and affects proper skeletal muscle trophism ^47,48^. MICU1-deficient mice also showed severe growth defects ^49^ and marked muscle weakness, easy fatigue and smaller muscle fibers ^49,50^. In addition, loss-of-function mutations in human *MICU1* are associated with myopathy ^51–53^. Together these observations corroborate the role of Tmbim5 in the regulation of growth and muscle function and point out similarities between the effects of Tmbim5, Mcu and Micu1 loss.

Two recent studies demonstrated that TMBIM5 functions as a pH-dependent mitochondrial Ca^2+^ efflux transporter ^28,29^. Zhang et al., in contrast, found that TMBIM5 overexpression in HEK cells enhanced mitochondrial Ca^2+^ uptake following ER Ca^2+^ release ^27^, suggesting that TMBIM5 may operate bidirectionally under certain conditions. To clarify the physiological relevance of this, we now generated two double knockout zebrafish lines: one lacking both Tmbim5 and Mcu, the primary mitochondrial Ca^2+^ influx mediator, and another deficient in Tmbim5 and Slc8b1, the zebrafish ortholog of NCLX, the presumed major mitochondrial Ca^2+^ efflux pathway. Surprisingly, both double knockout lines were viable and exhibited no overt organismal phenotypic alterations.

The absence of enhanced mitochondrial Ca^2+^ uptake deficits in *tmbim5*/*mcu* double knockout zebrafish argues against Tmbim5 functioning as a primary Ca^2+^ influx channel as this would have likely caused a more severe phenotype and would have reduced Ca^2+^ uptake independently of Mcu function. Two alternative mechanisms remain plausible to reconcile these data with previous reports: First, Tmbim5 may facilitate Ca^2+^ transport specifically at high Ca^2+^ concentrations. The increased uptake observed in TMBIM5-overexpressing cells after ATP stimulation ^27^ might be relevant at mitochondria-ER contact sites where local Ca^2+^ reaches up to 100 μM – conditions not tested in our system. Second, Tmbim5 may function as a regulatory modifier of MCU complex activity rather than as an autonomous Ca^2+^ influx channel. This hypothesis is supported by recent proteomic analyses identifying Tmbim5 as a component of the MCU interactome ^43^. The normal mitochondrial Ca^2+^ uptake observed in Tmbim5-deficient mitochondria also argues against significant alteration of MCU complex function through the proposed AFG3L2-EMRE regulatory axis ^54,55^. In contrast, while the reduced basal Ca²⁺ levels in *tmbim5*/*slc8b1* muscle could indicate decreased Ca^2+^ influx, we interpret this as evidence of severely compromised mitochondrial integrity.

Our study provides the first comprehensive characterization of zebrafish *slc8b1*, establishing its expression pattern, functional properties, and loss-of-function phenotype as the probable NCLX ortholog in this model organism. Similar to murine NCLX ^18^, zebrafish *slc8b1* demonstrates predominant expression in neural tissues. Functional analyses confirmed that Slc8b1 depletion significantly attenuated Na^+^-dependent mitochondrial Ca^2+^ efflux but residual Na^+^-dependent efflux persisted suggesting potential compensatory mechanisms or alternative transport pathways, possibly Tmem65 ^19–21^. Interestingly, in contrast to *slc8b1*, depletion of fish *tmem65* results in lethality ^56^. The structural divergence between zebrafish Slc8b1 and its mammalian counterparts – with fish Slc8b1 being approximately half the size of human NCLX – also raises intriguing mechanistic questions. It is possible that zebrafish Slc8b1 may function as a homodimer, as it was observed for mammalian NCLX ^57^, or potentially interact with additional protein partners to constitute a fully functional NCLX complex ^57^.

Notably, the effects of the loss of Slc8b1 in *tmbim5* knockout fish differed depending on the tissue. Also in mammals, *Nclx* knockout in cardiac tissue causes lethal mitochondrial calcium overload and depolarization ^18^. In neurons, *Nclx* knockout leads to mitochondrial calcium overload, which disrupts synaptic calcium transients and impairs synaptic plasticity, affecting learning and memory ^45,58,59^. In astrocytes, however, *Nclx* knockout increases glycolytic flux and lactate secretion, leading to improved cognitive performance ^45^. Overall, even the transport function of Nclx is not completely understood as *Slc8b1* is highly expressed in mouse liver tissue, where Na^+^-dependent Ca^2+^ efflux is minimal ^60^. The arrival of TMEM65 as an additional transport system or NCLX regulator or interacting partner ^19–21^ will hopefully bring some more clarity.

The viability of *tmbim5/slc8b1* double knockout fish, despite compromised mitochondrial Ca²⁺ handling, suggests the presence of robust compensatory mechanisms or alternative pathways that maintain Ca²⁺ homeostasis when both major efflux systems are disrupted. LETM1 represents a prime candidate for such compensation. LETM1 has been proposed to mediate mitochondrial Ca²⁺/H⁺ exchange ^15^, though it was originally identified as a component of the mitochondrial K⁺/H⁺ exchanger ^16^. While LETM1-containing liposomes demonstrated Ca²⁺ transport capacity ^15,61,62^, its physiological role as a direct Ca²⁺ transporter remains disputed.

Interestingly, LETM1 might interact with TMBIM5 ^28^, suggesting potential functional crosstalk between these proteins. In zebrafish, *letm1* deficiency causes embryonic lethality in most but not all mutants, suggesting variable compensatory capacity ^63^. However, the specific role of Letm1 in Ca²⁺ homeostasis has not been functionally characterized in this model.

Intriguingly, our gene expression analysis revealed tissue-specific regulation of *letm2*, a LETM1 paralog, rather than *letm1* itself. In *tmbim5/slc8b1* double knockouts, *letm2* was downregulated in brain tissue (where phenotypes were ameliorated) but upregulated in muscle tissue (where dysfunction was exacerbated). If Letm2 were functioning compensatorily, one would expect the opposite pattern. Unlike LETM1, LETM2 lacks Ca²⁺-binding EF-hand domains and cannot functionally replace LETM1, as evidenced by its inability to rescue LETM1 knockdown-induced mitochondrial swelling^44^). One speculative explanation could be that LETM2 functions as a negative regulator of LETM1 activity, analogous to how MCUb inhibits MCU-mediated Ca²⁺ uptake ^64^, though this hypothesis requires experimental validation and remains highly tentative given the limited functional data available for LETM2. Alternatively, the reduced mitochondrial mass might trigger a beneficial shift in energy metabolism towards glycolysis, similar to the beneficial metabolic adaptation observed in astrocyte-specific *Nclx* knockout mice ^45^.

In conclusion, loss of Tmbim5 impairs zebrafish growth, increases cell death in the brain and results in abnormal mitochondrial function as evidenced by reduced mitochondrial membrane potential. However, the absence of Tmbim5 did not exacerbate the phenotype of either *mcu* or *slc8b1* knockout zebrafish despite functional and morphological changes in *tmbim5*/*slc8b1* double knockout fish. The effects of Slc8b1 loss in Tmbim5 knockout zebrafish are tissue-specific. We demonstrated that Slc8b1 depletion can either restore or exacerbate the Tmbim5 absence phenotype.

## Materials and methods

### Animal Maintenance

*tmbim5^+/+^, tmbim5^−/−^*, *tmbim5*^+/+^;*mcu*^-/-^, *tmbim5*^-/-^;*mcu*^-/-^, *tmbim5*^+/+^;*slc8b1*^-/-^ and *tmbim5*^-/-^;*slc8b1*^-/-^ zebrafish were used in the study. All animals were maintained following previously described methods ^67^. in the Zebrafish Core Facility, a licensed breeding and research facility (PL14656251, registry of the District Veterinary Inspectorate in Warsaw; 064 and 051, registry of the Ministry of Science and Higher Education) at the International Institute of Molecular and Cell Biology in Warsaw. Adult zebrafish and larvae were kept in E3 medium (2.48 mM NaCl, 0.09 mM KCl, 0.164 mM CaCl_2_·2H_2_O, and 0.428 mM MgCl_2_·6H_2_O) at 28.5 °C. Larvae were kept in a Petri dishes (∼50 larvae/dish) in an incubator under a 14 h/10 h light/dark cycle. The stages of fish development were defined as hours postfertilization (hpf) and days postfertilization (dpf). Zebrafish embryos intended for stainings or *in vivo* imaging experiments, were treated with N-phenylthiourea (PTU, Alfa Aesar Cat# 103-85-5) at final concentration of 0.003% to avoid pigmentation.

This study follows ARRIVE guidelines ^68^. All experimental procedures were approved by the Local Ethical Committee for Experiments on Animals in Warsaw (permission no. WAW2/050/2022) and were performed in accordance with European and Polish regulations on animal welfare.

### Mutant and transgenic zebrafish lines

The generation of *tmbim5*^−/−^ zebrafish using CRISPR-Cas9 gene editing was previously described in Gasanov ^33^. Briefly, a mix of two gRNA: 5’-GGCTGGTATGTGGGAGTCTA-3’ and 5’-CGCGGGCAGTGTGGGCCTGA-3’ (0.04 mg/ml of each) and Cas9 protein (0.2 mg/ml) was injected into the cytoplasm of one-cell stage wild-type AB zebrafish embryos.

Fish were raised after injections up to 4-months-old and were fin-clipped for DNA sampling and high resolution melting (HRM) analysis. Targeted region in the 4^th^ exon of *tmbim5* was amplified using Precision Melt Supermix (BioRad, Cat# 1725112), primers: *tmbim5 HRM forward*: 5’-CGATCTGGCCGCAGTACG-3’, *tmbim5 HRM reverse*: 5’-CCAGCCAGGAGTTGCTC-3’ and the CFX Connect RT-PCR Detection System (BioRad). Melting curve analysis was performed using Precision Melt Analysis (Bio-Rad) software. The mutation-positive ones were analyzed by DNA sequencing. Fish with the deletion of 29 nucleotides and premature STOP codon was selected as F0 founder. F0 fish was outcrossed with the AB zebrafish line, and their mutation-carrying offspring were in-crossed to generate homozygous mutants. Fish were screened via PCR using the same primers as for HRM analysis, followed by Sanger sequencing.

Genotyping of subsequent generations of *tmbim5*^+/+^, *tmbim5*^+/-^ and *tmbim5*^-/-^ fish was based on the difference in the size of PCR product that was detected by electrophoresis in 2% agarose.

To generate double knockout of *tmbim5* and *mcu*, *tmbim5*^-/-^ zebrafish were outcrossed with *mcu*^-/-^ zebrafish (obtained as a kind gift from prof. Jacek Kuźnicki laboratory). Obtained heterozygous fish were incrossed and their offspring was genotyped for *tmbim5* as described above and for *mcu* as described in Soman et al.^11^_._

To generate zebrafish knockout of *nclx* the exon 5 of *slc8b1* gene was targeted with gRNA designed with free web tools: CHOPCHOP ^65^ and CRISPRscan ^66^ according to the protocol described in Wiweger et al. ^69^. Sequences that were common in both of the predictions and were scored as low risk for off-targets were chosen. The correctness of the sequence in the targeted area in the AB zebrafish was confirmed by Sanger sequencing. The gene-specific oligonucleotide with T7 overlaps: 5’-taatacgactcactataGGAGTGACGTTCCTGGCTCTgttttagagctagaa-3’. During gRNA preparation, annealing and filling in steps were combined, and the template was prepared by PCR using BioMix Red (BioLine, Cat# BIO-25006), gene-specific and constant oligo: 5’-AAAAGCACCGACTCGGTGCCACTTTTTCAAGTTGAT AACGGA CTAGCCTTATTTTAACTTGCTATTTCTAGCTCTAAAAC-3’ (10 µM). The PCR conditions were the following: pre-incubation at 95°C for 3 min, followed by 35 cycles of 95°C for 15 s, 40°C for 15 s, and 68°C for 15 s. sgRNA was prepared using 150 ng of purified PCR product and MegaScript T7 Transcription Kit (Invitrogen, Cat# A57622) according to the manufacturer’s recommendations. Thereafter, RNA was precipitated by overnight incubation with ammonium acetate at -20°C.

gRNA was suspended in water, and the concentration was adjusted to 500 ng/μl. Cas9 protein (14 mg/ml stock, made in-house) was diluted in KCl/HEPES (200 mM/10 mM, pH 7.5) buffer to a final concentration of 600 ng/μl. The injection mixture was assembled fresh by mixing 2 μl of gRNA (1 μg), 2 μl of Cas9 (1.2 μg), and 0.5 μl phenol red (Sigma, Cat# P0290) and left for complex formation at room temperature for 5–10 min. Thereafter, the samples were kept on ice. Microneedles were pulled from borosilicate glass capillaries (Sutter; Cat# B100-75-10) using a P-1000 Flaming/Brown micropipette puller (Sutter). One nanoliter of the gRNA/Cas9/phenol red mixture was injected into the yolk at 1–2 cell-stage *tmbim5*^-/-^ zebrafish embryos using a FemtoJet microinjector (Eppendorf).

Fish were grown up after injections up to 4-months-old and were fin-clipped for DNA sampling and high resolution melting (HRM) analysis. Targeted region in the 5^th^ exon of *scl8b1* was amplified using Precision Melt Supermix (BioRad, Cat# 1725112), primers: *slc8b1 HRM forward*: 5’-GTGTACCATGTGGATGTGTGTTG-3’, *slc8b1 HRM reverse*: 5’-CAGACCAGCAGTTTGAGGGTG-3’ and the CFX Connect RT-PCR Detection System (BioRad). Melting curve analysis was performed using Precision Melt Analysis (Bio-Rad) software. The mutation-positive ones were analyzed by DNA sequencing of PCR products. Samples for sequencing were prepared by PCR using BioMix Red (BioLine, Cat# BIO-25006), *slc8b1 seq forward*: 5’-GAGTTTGCCGTTTA ATTTCTGG-3’, *slc8b1 seq reverse*: 5’-AATTCCTCACCAAAAAGTGCTC-3’ and 2 μl of gDNA template per 20 μl of reaction. Fish with the confirmed insertions or deletions leading to premature STOP codon were selected as F0 founders. F0 fish was outcrossed with the AB zebrafish line, and their mutation-carrying offspring were in-crossed to generate double-homozygous mutants. Fish were screened via HRM analysis, followed by Sanger sequencing.

### Transient expression of 4mtD3cpv Ca^2+^ probe

To assess basal mitochondrial Ca^2+^ levels in the brain in skeletal muscle of zebrafish embryos *in vivo*, the genetically encoded FRET-based Ca²⁺ probe 4mtD3cpv was used. The probe was expressed by microinjection of plasmid DNA encoding 4mtD3cpv under the control of the human cytomegalovirus (CMV) promoter ^38^ (a kind gift from the laboratory of Prof. Wolfgang F. Graier). Approximately 1 nl of plasmid DNA solution (400 ng/µl) was injected into the cell of one-cell stage embryos to obtain transient and mosaic expression of the probe. Embryos were screened for expression at 1 dpf using fluorescence microscopy. Only those showing appropriate mitochondrial localization of the probe were selected for imaging. Microinjections were performed as described above.

### Quantitative Real-Time PCR

Total RNA was extracted from pools of 15–20 larvae or dissected tissues dissected from two adult zebrafish using TRI Reagent (Invitrogen, Cat# AM9738) according to a previously published protocol ^70^. RNA concentration and purity were measured using a NanoDrop spectrophotometer. Only samples with absorbance >1.8 at A260/280 nm were used for further processing. Reverse transcription was performed with 500 ng of total RNA using iScript™ cDNA Synthesis Kit (BioRad, Cat# 1708891).

Quantitative PCR was conducted using SsoAdvanced Universal SYBR Green Supermix (BioRad, Cat# 1725274) on a CFX Connect RT-PCR Detection System (Bio-Rad). Primer sequences are listed in Table S1. Relative gene expression levels were calculated using the 2^ΔΔCq^ method, with *rpl13a*, *ef1a*, or *18S* used as reference genes depending on tissue type. Each reaction was performed in technical duplicate using 25 ng of cDNA and 0.25 μM of each primer. Changes in expression are expressed as fold changes using expression in WT as a reference value. The calculations were performed in Microsoft Excel.

### Quantification of mitochondrial (mtDNA) to nuclear DNA (nDNA) ratio

DNA was isolated from zebrafish following a previously published protocol (Otten *et al*, 2020). Adult zebrafish were euthanized with tricaine (MS-222) and sacrificed to obtain a single DNA sample. The samples were incubated in a lysis buffer (75 mM NaCl, 50 mM EDTA, 20 mM HEPES pH 7.5, 0.4% SDS, 1 mg/ml proteinase K) for at least 4 h in 55°C. For adult zebrafish, brain and skeletal muscle samples were dissected and homogenized with a plastic pestle in lysis buffer before incubation. DNA was precipitated with isopropanol and incubated for at least 6 h in -20°C. Following centrifugation, the DNA pellet was washed with 70% ethanol, air-dried, and dissolved in Tris-EDTA (TE) buffer. Relative mitochondrial DNA (mtDNA) abundance was quantified by measuring the levels of *mitochondrial NADH dehydrogenase* (*mt-nd1*) and *nuclear elongation factor 1-alpha* (*ef1a*) using qRT-PCR (primers listed in Table 1). A total of 25 ng of DNA was used per qRT-PCR reaction. Changes in the mtDNA-to-nDNA ratio were calculated as fold changes using WT as a reference. Calculations were performed in Microsoft Excel.

### Whole-mount *in situ* hybridization

Whole-mount in situ hybridization (WISH) was performed on embryos fixed overnight in 4% paraformaldehyde (PFA) at room temperature. Fixed embryos were dehydrated in a graded methanol series and stored at -20°C in 100% methanol. Whole-mount in situ hybridization (WISH) was performed as previously described (Soman et al. 2019) with several modifications. Prior to hybridization, embryos were rehydrated through a methanol/PBST (PBS + 0.1% Tween-20) series and treated with 0.1% sodium citrate and 0.1% Triton X-100 for 10 min on ice for permeabilization. Pre-hybridization was performed in Hybridization Solution (HS; 50% deionized formamide, 5X sodium citrate buffer (SSC), 500 μg/ml tRNA, 50 μg/ml heparin, 0.1% Tween-20, 10mM citric acid, 5% sodium dextran sulfate) at 65°C for 3 h. DIG-labeled antisense RNA probes were synthesized by *in vitro* transcription using MegaScript T7 Transcription Kit (Invitrogen, Cat# A57622), digoxigenin (DIG)-labeled UTP (Roche, Cat# 11277073910) and template DNA generated by PCR amplification of target gene sequences. Hybridization was carried out at 67°C overnight in the hybridization buffer (0.5-3 μg probe/ml buffer). After stringent washes with Hybridization Wash Solution (HWS; 50% deionized formamide, 5X SSC, 0.1% Tween-20, 10 mM citric acid) and subsequently with 75% HWS, 50% HWS, 25% HWS in 2X SSCT (SSC with 0.1% Tween-20) and 0.2X SSCT at 65°C, embryos were blocked with 10% Blocking Reagent (Roche, Cat# 11096176001) in Maleic Acid buffer (100 mM maleic acid, 100 mM NaCl, 50 mM MgCl_2_, 0.1% Tween-20) and incubated with an alkaline phosphatase-conjugated anti-DIG antibody (1:5000, Roche, Cat# 11093274910) overnight at 4°C. Staining was developed using nitro-blue tetrazolium chloride (0.45 mg/ml, Roche, Cat# 11383213001) and 5-bromo-4-chloro-3-indolyl-phosphate (0.175 mg/ml, Roche, Cat# 11383221001) in Detection buffer (50 mM Tris-HCl pH 9.5, 50 mM NaCl, 25 mM MgCl_2_, 0.05% Tween 20, 2% polyvinyl alcohol) for 3 h and monitored under a stereomicroscope. Background signal was cleaned by 10 min wash with 100% methanol and samples were re-fixed with 4% PFA for 20 min at room temperature and stored in the glycerol until imaging. Images were captured with a Nikon SMZ25 stereomicroscope.

### Histology

For histological analyses, eight-month-old zebrafish were fixed in Bouin’s solution, dehydrated in ethanol, xylene and embedded in paraffin using standard protocols. Embedded samples were sectioned at 6 µm thickness using a Leica microtome (RM2265, Leica). Sections were stained with Hematoxylin and Eosin (H&E) for general tissue morphology, Alcian Blue–Periodic Acid Schiff (AB-PAS) for mucopolysaccharides and glycogen, or Masson’s Trichrome for collagen visualization. Slides were imaged using a Nikon Eclipse 90i microscope with a Nikon DS5-U1 camera (Nikon Corporation, Japan) and the computer image analysis system NIS-Elements AR (Nikon Corporation, Japan). The quantification of Alcian Blue and Masson’s trichrome staining intensity and area was performed using the Colour Deconvolution plug-in in ImageJ.

### Isolation zebrafish larvae mitochondria

Mitochondria were isolated from zebrafish larvae at 5 dpf as previously described by Prudent et al. ^71^, with slight modifications. Approximately 300-450 larvae were pooled per biological replicate. Larvae were euthanized with Tricaine and homogenized in 4 ml of cold isolation buffer (210 mM mannitol, 70 mM sucrose, 1mM EDTA, 10mM HEPES, pH 7.5, supplemented with 2 mM PMSF and 2 mg/ml BSA) using a glass-Teflon homogenizer. Homogenates were centrifuged twice at 1,500 g for 10 min at 4°C to remove debris and nuclei. The supernatant was transferred to a new tube and centrifuged at 14,000 g for 15 min at 4°C. The mitochondrial pellet was washed twice in an ice-cold KCl medium (125 mM KCl, 2 mM K_2_HPO_4_, 1 mM MgCl_2_, 5 mM glutamate, 5 mM malic acid, 20 mM HEPES, pH 7) and resuspended in the same buffer. Protein concentration was measured using the Bradford assay (BioRad, Cat# 5000006). Mitochondrial suspensions were kept on ice and used within 1 h after preparation.

### Ca^2+^ measurements in isolated mitochondria

Mitochondrial Ca^2+^ fluxes were measured in freshly isolated mitochondria from 5 dpf zebrafish larvae. 25 µg of mitochondria were gently suspended in KCl medium with glutamate (5 mM), malic acid (5 mM) freshly added, and transferred into a 96-well half-area black polystyrene microplate (Corning®, Cat# CLS3694). Thapsigargin (0.2 µM, Invitrogen™, Cat# T7459) was added in order to inhibit SERCA pump from potential contaminants from the endoplasmic reticulum in the samples. The bath [Ca^2+^] was detected using the Ca^2+^-sensitive dye Oregon Green™ 488 BAPTA-5N hexapotassium salt (2 µM, Invitrogen™, Cat# O6812) as described in Soman et al.^11^. The signal was measured using a plate reader (Tecan Infinite M1000 Pro) at excitation 494 nm/emission 521 nm. Measurements were performed at room temperature.

For the analysis, the first readout of baseline recording was defined as F_0_ and used for normalization. All traces are shown as ΔF/F_0_, where ΔF = F - F_0_.

Mitochondrial Ca^2+^ uptake assay: 20 µM CaCl_2_ diluted in KCl medium wasThe signal was measured every 3 s for 10 min. Ca^2+^ uptake was calculated as the difference between the maximal ΔF/F_0_ (first read-out after CaCl_2_ addition) and ΔF/F_0_ value measured 2 min after CaCl_2_ addition.

Na^+^-dependent mitochondrial Ca^2+^ efflux: Mitochondria were first loaded with 20 µM CaCl_2_ diluted in KCl medium, as described for the mitochondrial uptake assay. Next, mitochondrial Ca^2+^ uptake via MCU was inhibited by adding 10 µM Ru360 (Sigma-Aldrich, Cat# 557440). To estimate the amount of Na^+^-independent Ca^2+^ efflux following Ru360 addition, the last ΔF/F_0_ value recorded after CaCl_2_ addition was subtracted from the maximum ΔF/F_0_ value obtained after Ru360 addition. Subsequently, Na^+^-dependent mitochondrial Ca^2+^ efflux was stimulated by adding 10 mM NaCl (KCl medium in which 120 mM K^+^ was substituted with Na^+^). To estimate the amount of Ca^2+^ efflux following Na^+^ addition, the last ΔF/F_0_ value recorded after Ru360 addition was subtracted from the maximum ΔF/F_0_ value after Na⁺ addition. Fluorescence signals were recorded every 5 seconds for 5 minutes following each treatment.

### TMRE microinjections

The stock solution of tetramethylrhodamine ethyl ester (TMRE; Invitrogen™, Cat# T669) was prepared by dissolving in dimethyl sulfoxide (DMSO, Sigma-Aldrich, Cat# D8418) to a final concentration of 10 mM. The injection solution was prepared at the day of the experiment by diluting the 1:5 stock solution in water (final concentration: 2 mM) and 1 nl was injected into the yolk of embryos at the 1-2 cell stage. Microinjections were performed in the same way as for the generation of *nclx*^-/-^ mutants. Embryos were then grown according to standard methods described in the “Animal maintenance” section.

### *In vivo* imaging with LSFM

Stained or transgenic larvae were anesthetized with 0.2 mg/ml Tricaine (Sigma, Cat# A-5040), mounted in 2% low-melting-point agarose (Sigma-Aldrich, Cat# A9414) in a glass capillary and imaged under a Zeiss Lightsheet Z.1 microscope using a 40X objective (Zeiss) with excitation at 488 nm (AO), 561 nm (TMRE) or 445 nm (4mtD3cpv). Fish were imaged from the dorsal side of the head and were pushed from the capillary to be submerged in the medium. Measurements were performed at room temperature.

Images were first processed in ZEN software (Zeiss) in order to obtain merged maximum projection images, exported to TIF files and analyzed further with ImageJ.

Cell death estimation by Acridine Orange *in vivo* staining: Acridine Orange (AO; Sigma-Aldrich, Cat# A8097) was added to the plates containing live 4 dpf zebrafish larvae (final concentration: 10 µg/ml in E3). After 1 hour of incubation in the dark, the medium was exchanged with fresh E3 six times during a period of 30 min. Z-stacks encompassing 45 µm of the optic tectum with 5 µm interval were acquired with a Zeiss Lightsheet Z.1 microscope. The counting of the AO-stained cells in the optic tectum of each larvae, which were considered as dying ^72^, was performed manually on the maximum projection images with the CellCounter plug-in in ImageJ. The number of cells was normalized to the analyzed area of the optic tectum.

Basal mitochondrial Ca^2+^ measured *in vivo* with 4mtD3cpv: the use of the FRET-based Cameleon 4mtD3cpv allowed for ratiometric imaging and enabled estimation of basal Ca^2+^ levels, independent of probe expression level ^39,40^. Brains and somites of 2 dpf dechorionated larvae injected with the pcDNA3-4mtD3cpv vector were imaged using a Zeiss Lightsheet Z.1 microscope. The intensities of cyan fluorescent protein (CFP, emission: 460-500 nm) and circularly permuted Venus (cpV, emission: 522-565 nm) were recorded every 12 s for 1 min in a z-stack of 10 µm with 1 µm intervals.

During image analysis, the StackReg plug-in was used for motion correction. Cells in the skeletal muscle cells of the optic tectum that showed 4mtD3cpv expression were manually selected as ROIs. Mitochondria were detected by thresholding, and mean CFP and cpV fluorescence were quantified in each ROI. The mean ratio of cpV to CFP was calculated for the brain and muscles of each larva.

Mitochondrial membrane potential (MMP) *in vivo* measurements: Two days after microinjections with TMRE dechorionated larvae were imaged using lightsheet fluorescence microscopy (LSFM) as described above. 2 dpf larvae were treated with 10 μM CCCP (Sigma-Aldrich, Cat# C2759) to induce depolarization of mitochondria. To gain complete depolarization, larvae were treated for 5 min. Z-stacks encompassing 10 µm of the optic tectum with 1 µm intervals were acquired with a Zeiss Lightsheet Z.1 microscope every 5 s during 5 min of treatment. F was normalized to the value obtained after full depolarization (F/F_CCCP_) and the ratio between basal fluorescence (F_0_) and F_CCCP_ was calculated to compare mitochondrial membrane potential between variants. Image analysis was performed as for mitochondrial Ca^2+^ measurements described above.

### Coiling activity of zebrafish embryos

Randomly selected embryos were transferred to a 12-well plate (4 embryos/well) in 2 ml of E3 and acclimated for 2h in 28°C. Coiling activity was recorded at 30 hpf using the Nikon SMZ25 stereomicroscope for 3 min with a 10 frames /s acquisition rate. The number of coiling events for each embryo was counted manually.

### Transmission electron microscopy

Zebrafish larvae at 5 dpf were euthanized with Tricaine. The animals were fixed in 2.5% glutaraldehyde for 24 hours at 4°C, washed in PBS, postfixed in 1% osmium tetroxide for 1 hour, washed with water, and stained with 1% aqueous uranyl acetate overnight at 4°C. The larvae were dehydrated and infiltrated with epoxy resin (Sigma Aldrich, Cat# 45-359-1EA-F). Samples were then polymerized at 60°C for 48 hours.

Polymerized blocks were trimmed with a tissue processor (Leica EM TP) and cut with an ultramicrotome (EM UC7, Leica) to obtain ultrathin sections (70 nm thick), which were collected on nickel grids (200 mesh, Agar Scientific, Cat# G2200N). The grids were examined using a Tecnai T12 BioTwin transmission electron microscope (FEI) equipped with a 16-megapixel TemCam-F416 camera (TVIPS GmbH) at the Microscopy and Cytometry Facility at IIMCB in Warsaw.

### Quantification and statistical analysis

All statistical analyses were conducted using R version 4.1.2. Box and whisker plots were generated with R, while other graphs were generated with Microsoft Excel or GraphPad Prism version 8. All datasets were tested for outliers using a Grubbs’ test and for normal distribution using the Shapiro-Wilk test. The applied statistical tests are indicated in the figure legends.

### Data availability

This study includes no data deposited in external repositories. No unbiased larger data sets were generated in this study. All data are presented in the results or as supplemental data. All unique zebrafish lines generated in the study are available with a completed materials transfer agreement. Any additional information required to reanalyze the data reported in this paper is available upon request.

## Acknowledgments

We thank the Laboratory of Neurodegeneration at IIMCB and specifically Prof. Jacek Kuznicki for providing laboratory space to conduct research on zebrafish, mcu^-/-^ zebrafish line and helpful discussions. We thank Prof. Jacek Kuźnicki and Prof. Barbara Zabłocka for critical reading of the paper. We acknowledge the IIMCB Zebrafish Core Facility for service and fish material and Microscopy and Cytometry Facility for providing access to LSFM. This research was funded by the National Science Centre, Poland (021/40/C/NZ4/00031) to I.W.

## Author contributions

I.W. designed and performed most of the experiments with contributions from S.B. and Ł.M., D.A-U. performed histological analysis of adult zebrafish, M.M. and A.S. performed TEM analysis of larvae mitochondria, A.M. supervised experiments, I.W analyzed data and interpreted results, I.W. and A.M. wrote the manuscript.

## Supplemental material

### Tables

**Table S1.**
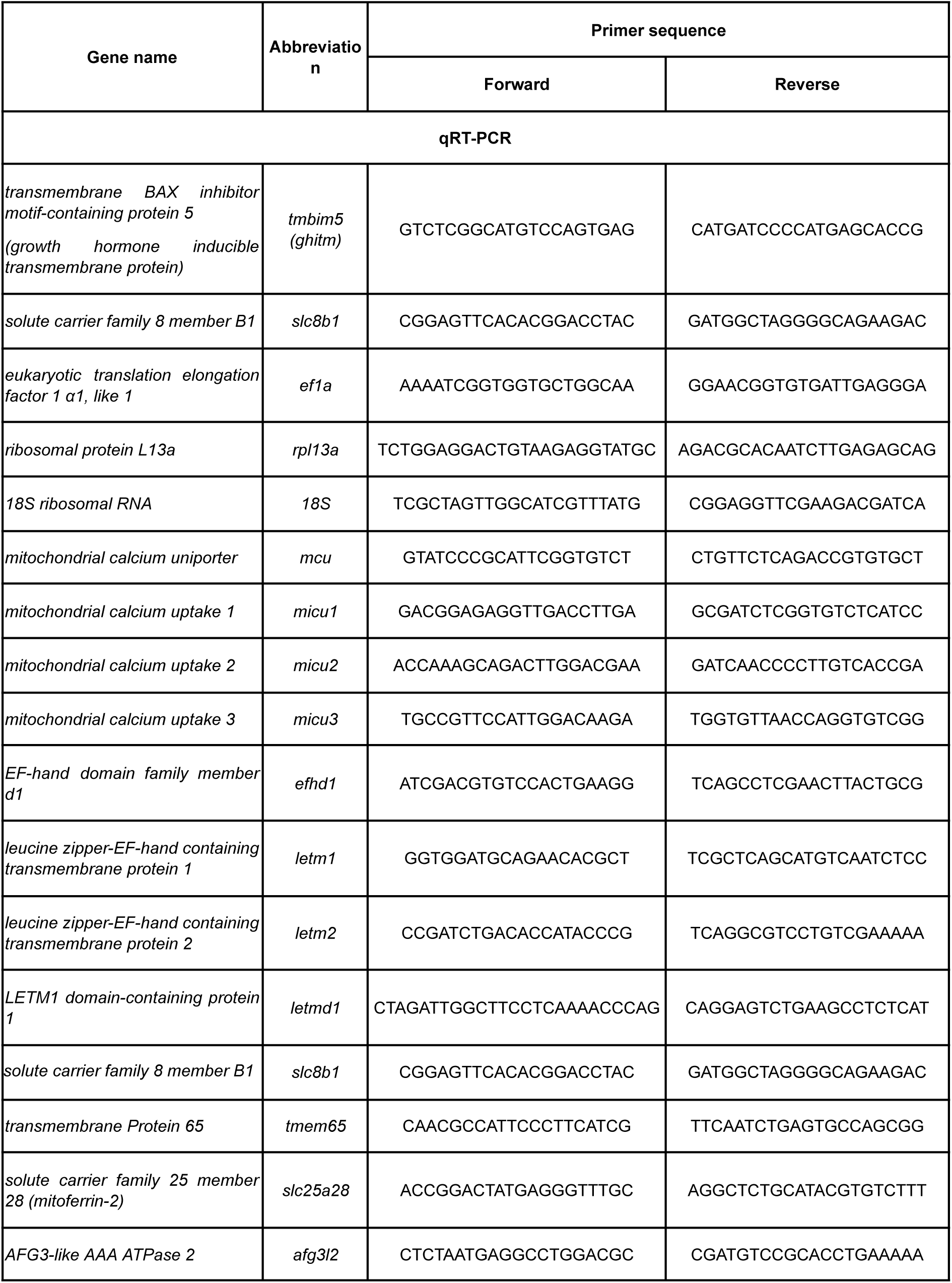
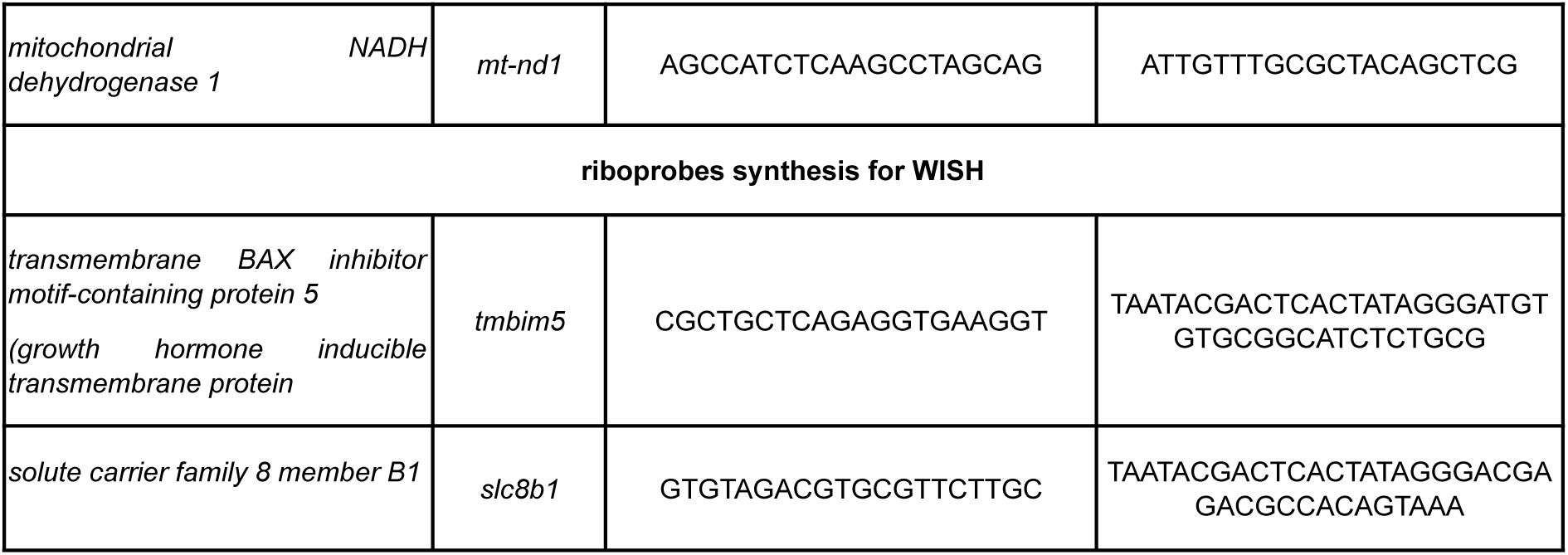
Primers that were used for riboprobe synthesis and qRT-PCR. All primers’ sequences are listed as from 5’ to 3’.

### Normal behavior of *tmbim5*^-/-^ zebrafish

**Figure S1.**
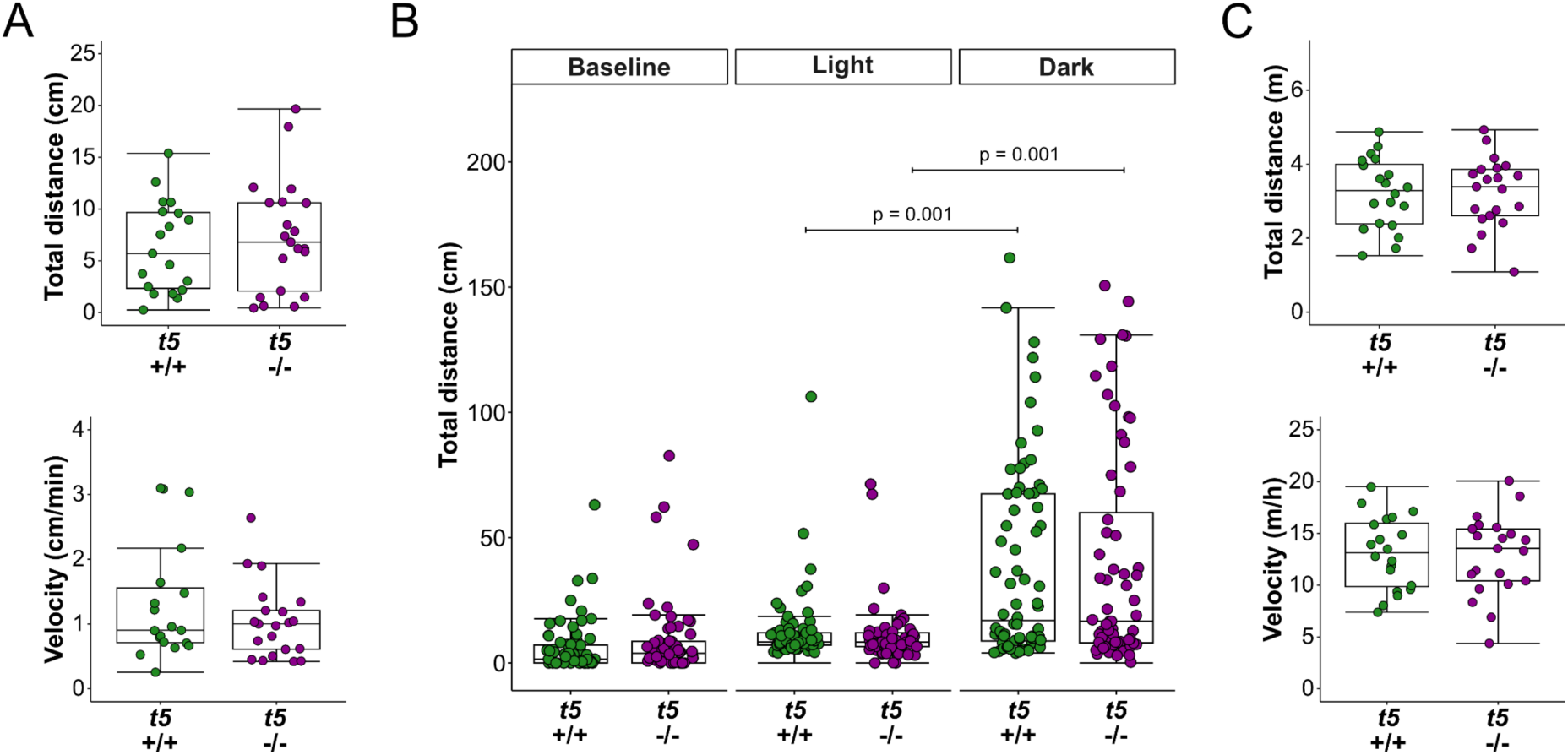
Normal behavior of *tmbim5*^-/-^ zebrafish. A) Locomotor activity of 5 dpf larvae in an open-field test is not affected by *tmbim5* knockout (KO). Total distance traveled and mean velocity are plotted as box-and-whisker plots (box: 25th–75th percentile; whiskers: min to max), with each dot representing an individual larva (*n* = 19–21). Number of experiments = 3. Statistical analysis: *t*-test (total distance) or Mann–Whitney test (velocity). B) Normal visual-motor response of 5 dpf *tmbim5*^-/-^ larvae. Total distance covered during each phase of the experiment is plotted as box-and-whisker plots, with each dot representing an individual larva (*n* = 64–72). Number of experiments = 3. Statistical analysis: Kruskal-Wallis test followed by Dunn’s test. C) Locomotor activity of adult (8-month-old) fish in a novel tank test is not affected by *tmbim5* KO. Total distance traveled and mean velocity are plotted as box-and-whisker plots, with each dot representing an individual fish (*n* = 20–21). Number of experiments = 3. Statistical analysis: Mann–Whitney test.

### Histology of brain, muscle and liver of adult *tmbim5*^-/-^ fish

**Figure S2.**
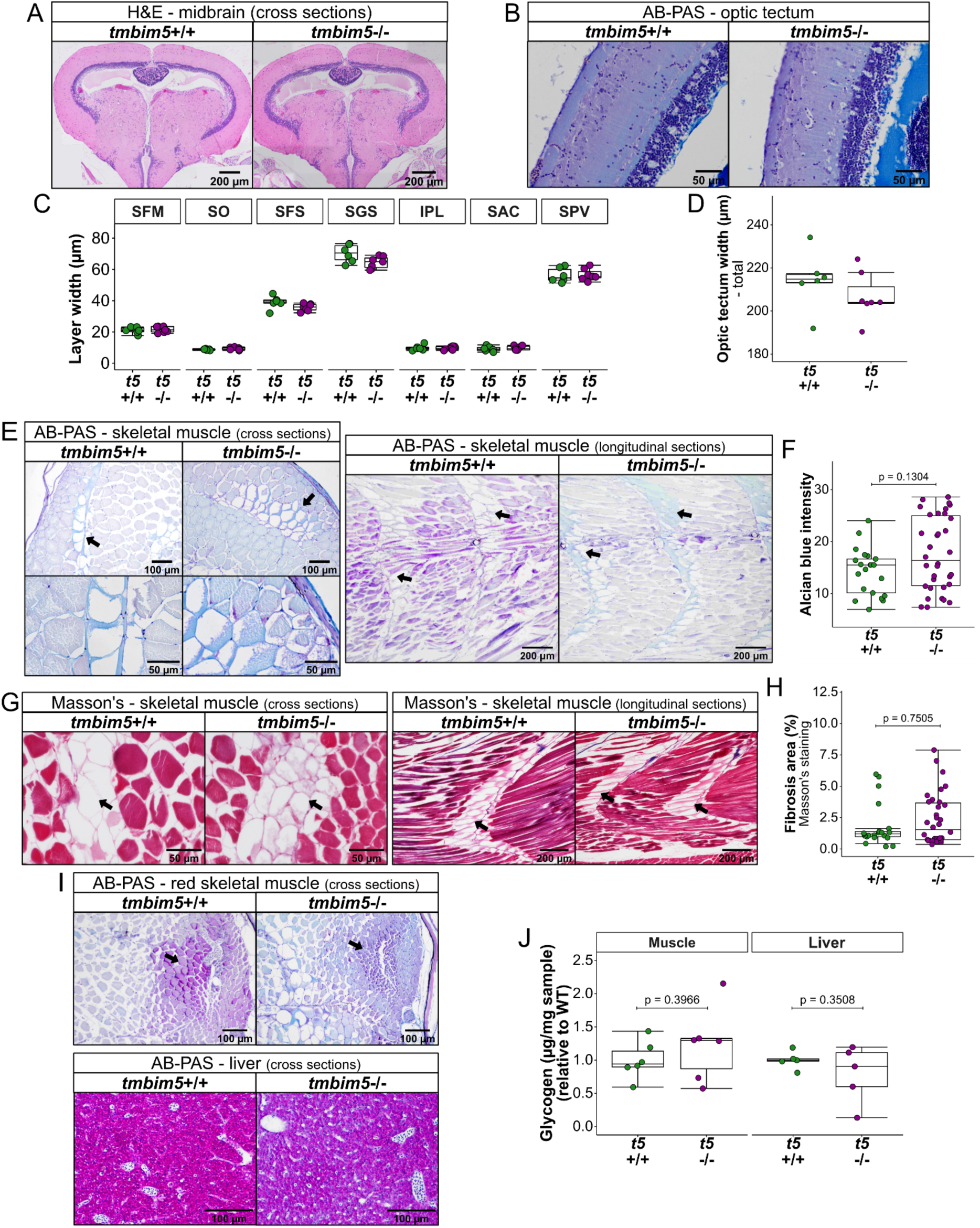
Histology of brain, muscle and liver of adult *tmbim5*^-/-^ fish. A-D) Normal histology of the *tmbim5*^-/-^ fish brain. A) Representative images of cross-sections of the WT and *tmbim5*^-/-^ midbrain stained with H&E. B) Representative images of cross-sections of the WT and *tmbim5*^-/-^ optic tectum stained with AB-PAS. C) Quantification of the width of optic tectum layers. D) Comparison of total optic tectum width between WT and *tmbim5*^-/-^ fish, showing no significant differences. Data are presented as box-and-whisker plots (box: 25th–75th percentile; whiskers: min to max), with each dot representing an individual fish (5–7 slices analyzed per fish, 5 measurements per slice, *n* = 6–7). Statistical analysis: *t*-test with BH correction for multiple comparisons. SFM – Stratum Fibrosum Superficiale, SO – Stratum Opticum, SFS – Stratum Fibrosum Superficiale, SGS – Stratum Fibrosum Griseum, IPL – Internal Plexiform Layer, SAC – Stratum Album Centrale, SPV – Stratum Periventriculare. E-F) Enhanced Alcian Blue (AB) staining in the intermuscular spaces of *tmbim5*^-/-^ fish. E) Representative images of cross- and longitudinal sections of WT and *tmbim5*^-/-^ fish stained with AB-PAS (Periodic Acid-Schiff). Lower panels show magnified views of AB-stained intermuscular spaces (indicated by arrows). F) Quantification of AB staining intensity. Data are presented as box-and-whisker plots, showing a trend toward increased staining intensity in *tmbim5*^-/-^ fish. Each dot represents the average for one slice (*n* = 21–34, number of fish = 7). Statistical analysis: Mann–Whitney test. G-H) Masson’s trichrome staining of skeletal muscle reveals no differences in *tmbim5*^-/-^ fish. G) Representative images of cross- and longitudinal sections of WT and *tmbim5*^-/-^ fish. Intermuscular spaces that showed positive AB staining are marked with arrows. H) Quantification of Masson’s trichrome-positive areas, showing no significant changes in *tmbim5*^-/-^ fish. Each dot represents the average for one slice (*n* = 21–34, number of fish = 7). Statistical analysis: Mann–Whitney test. I) Reduced PAS staining intensity in the muscle and liver of *tmbim5*^-/-^ adult (8-month-old) fish. Representative images of cross-sections of WT and *tmbim5*^-/-^ fish stained with AB-PAS. J) Glycogen levels remain unchanged in the muscle and liver of *tmbim5*^-/-^ adult (1.5-year-old) fish. Quantification was performed using a colorimetric glycogen assay. Each dot represents an independent biological replicate (*n* = 5–6, number of experiments = 2). Statistical analysis: *t*-test.

### Generation of Slc8b1-deficient fish

**Figure S3.**
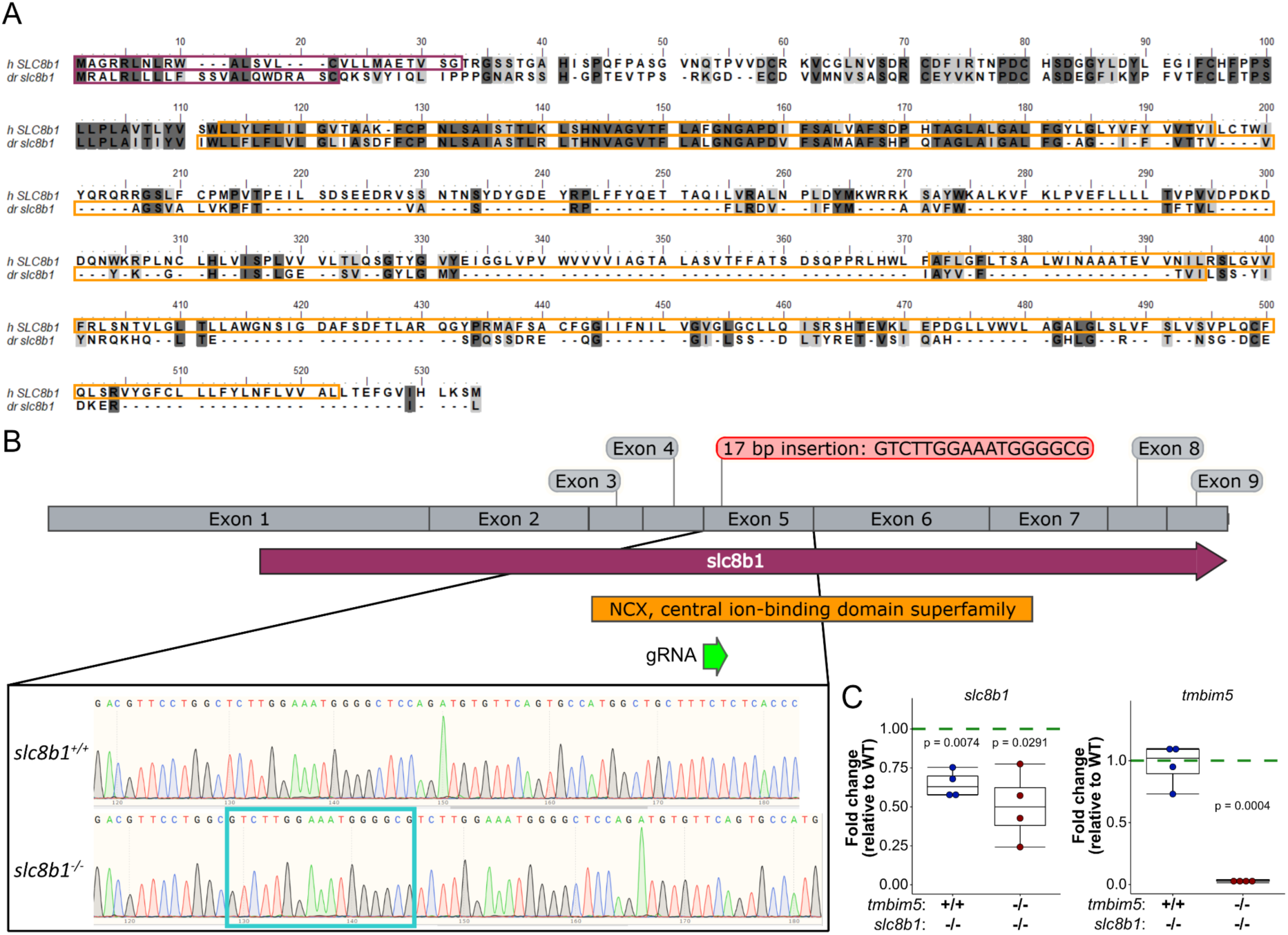
Generation of Slc8b1-deficient fish. A) Sequence alignment of human and zebrafish NCLX proteins encoded by *slc8b1*, performed using Optimal Global Alignment with the BLOSUM62 similarity matrix. The alignment revealed 24% sequence identity between species. Identical residues are highlighted in dark gray, while residues with similar physico-chemical properties are marked in light gray. The predicted signal peptide is enclosed within a violet frame, and the sodium/calcium (Na^+^/Ca^2+^) domain is highlighted by an orange frame. B) Schematic representation of the mutation site in *slc8b1*, accompanied by Sanger sequencing results. Inserted base pairs are highlighted with a turquoise frame. C) Expression levels of *slc8b1* and *tmbim5* mRNA in 5 dpf *tmbim5*^+/+^*;slc8b1*^-/-^ and *tmbim5*^-/-^*;slc8b1*^-/-^ larvae, quantified using qPCR and normalized to wild-type. *rpl13a* and *ef1a* were used as reference genes. Results are presented as box-and-whisker plots (box: 25th–75th percentile, whiskers: min–max), with each dot representing an independent biological replicate (*n* = 4, each RNA sample isolated from 30 larvae). Statistical analysis: one-sample *t*-test with BH correction for multiple comparisons.

### Normal behavior of *slc8b1*^-/-^ and *tmbim5*/*slc8b1* double knockout larvae

**Figure S4.**
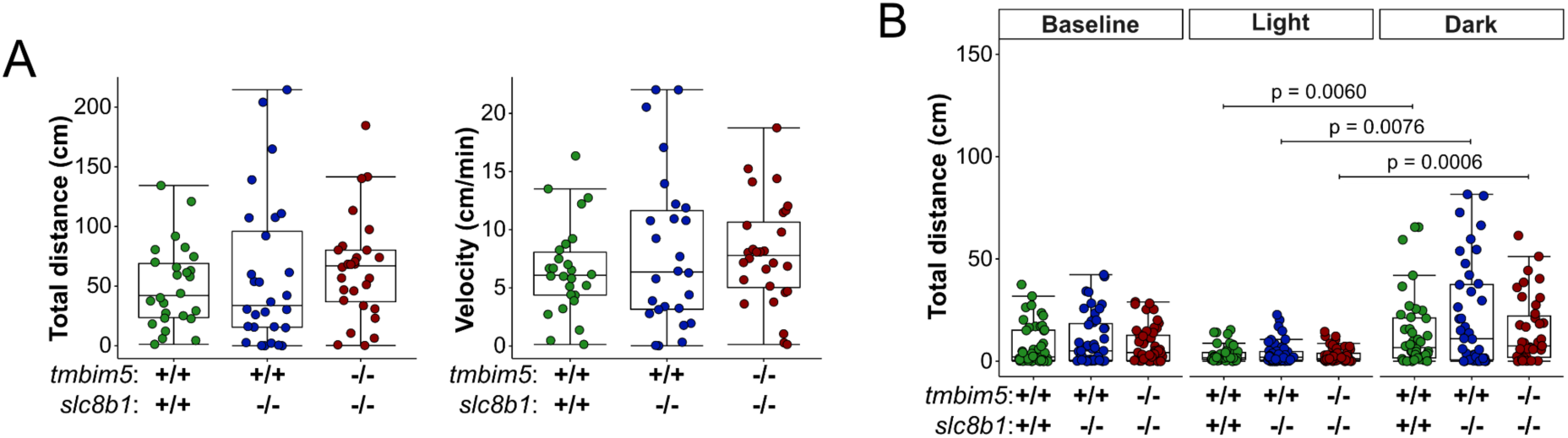
Normal locomotor activity of *slc8b1*^-/-^ and *tmbim5*/*slc8b1* double knockout larvae. A) Locomotor activity of 5 dpf larvae in an open-field test is not affected by *slc8b1* KO or *tmbim5*^-/-^*;slc8b1*^-/-^ double KO. Total distance traveled and mean velocity are plotted as box-and-whisker plots (box: 25th–75th percentile, whiskers: min–max), with each dot representing an individual larva (*n* = 26–28). Statistical analysis: Kruskal-Wallis test followed by Dunn’s post-hoc test. Number of experiments = 2. B) Normal visual-motor response of *slc8b1*^-/-^ and *tmbim5*^-/-^*;slc8b1*^-/-^ larvae at 5 dpf. Total distance traveled during each phase of the experiment is presented as box-and-whisker plots, with each dot representing an individual larva (*n* = 40–48). Statistical analysis: two-way ANOVA followed by Kruskal-Wallis test and Dunn’s post-hoc test. Number of experiments = 2.

### Supplemental Material and Methods

#### Glycogen levels quantification

Glycogen levels in the muscles and liver of adult (1.5-year-old) zebrafish were quantified using a colorimetric glycogen assay (Abcam, Cat# ab169558) according to the manufacturer’s instructions. Briefly, the assay is based on glycogen hydrolysis into glucose, which is oxidized to form an intermediate that reduces a colorless probe to a colored product with strong absorbance at 450 nm. Fish were euthanized with tricaine and dissected to obtain liver and muscle samples. Tissue was weighed before homogenization using a glass-Teflon homogenizer on ice. Glycogen levels were estimated by measuring the optical density (OD) at 450 nm using an absorbance microplate reader (Tecan Sunrise). Samples that were not treated with the glycogen-hydrolyzing enzyme served as background controls. Glycogen content was normalized to tissue weight and expressed as a fold change using WT as a reference.

#### Behavioral analysis

Open field test: Randomly selected 4 dpf larvae were acclimated to the behavioral testing room for at least 15 min. Two minutes before recording locomotor activity, the larvae were transferred to a 12-well plate that was then placed in the ZebraBox, a high-throughput monitoring system (ViewPoint). The experiment was performed in a volume of 2 mL of E3 medium, and the light intensity was set to 70%. Locomotor activity was recorded for 10 min.

Visual motor response: The experiment was performed according to the procedure described previously ^73^. On the day before the experiment, the larvae were placed in 24-well plates that contained 0.5 mL of E3 medium. Thirty minutes before recording locomotor activity, the plates were placed in the ZebraBox. The experiment consisted of three phases of the following changes in lighting conditions: baseline (0–10 min, 0% light intensity), light (10–20 min, 70% light intensity), and dark (20–30 min, 0% light intensity).

Novel tank test for adult zebrafish: The 8-month-old zebrafish were acclimated to the behavioral testing room for one week before the experiment. Fish were placed individually in a transparent tank (24 cm length x 14 cm height x 6cm width) filled with aquarium water. ZebraCube, an adult fish monitoring system (ViewPoint) was utilized to record animals’ behavior for 15 min.

The video files acquired during experiments with larvae and adult zebrafish were further analyzed using EthoVision XT software (Noldus).

